# Centrosome architecture and m6A-dependent gating of p53 surveillance after whole-genome doubling

**DOI:** 10.64898/2026.02.25.707964

**Authors:** Daniele Migliorati, Alessia Mattivi, Gian Mario Moretta, Selene Tessadri, Nicole Cona, Giorgia Pellizzaro, Michael Pancher, Mattia Furlan, Lucia Coscujuela, Martin Wegner, Marine H. Laporte, Michela Libergoli, Feryel Soualmia, Stefano Biressi, Toma Tebaldi, Alessandro Quattrone, Alberto Inga, Etienne D. Jacotot, Virginie Hamel, Paul Guichard, Mattia Pelizzola, Manuel Kaulich, Matteo Burigotto, Luca L. Fava

## Abstract

Whole-genome doubling (WGD) generates a tetraploid state that challenges cellular homeostasis and is strongly selected against in *TP53*-proficient contexts. Although centrosome amplification serves as a structural correlate of genome doubling, how centrosomal features engage and sustain p53 surveillance remains incompletely understood. Here, we develop a genetically encoded MDM2-based fluorescent reporter to monitor Caspase-2 pathway activation in living cells after WGD. Combining pharmacological profiling and genome-wide CRISPR screening, we uncover a multi-layered regulation of p53 responses. We identify PLK1-dependent centriole maturation and subdistal appendage integrity as architectural determinants of PIDDosome-dependent Caspase-2 activation, independent of distal appendage assembly. We show that upon WGD, Caspase-2-mediated cleavage of MDM2 reshapes the p53-MDM2 regulatory loop, enabling sustained p53 signaling. This sustained output depends on the catalytic activity of METTL3 and associated m6A writer proteins. Together, these findings demonstrate that centrosome architecture and epitranscriptomic regulation cooperate to shape p53 surveillance after genome doubling.

**Highlights:**

- Whole-genome doubling creates a surveillance state sensed by centrosome cluster architecture.
- Caspase-2 activation converts the p53-MDM2 loop from negative to positive feedback.
- m6A methylation establishes competence of the p53 module for sustained output.
- A genetically encoded biosensor enables functional dissection of this circuit.

## Introduction

Whole-genome doubling (WGD) generates a tetraploid state with profound consequences for genome stability^1-3^. While WGD can occur physiologically in specialized cell types through programmed polyploidization, such as in mammalian hepatocytes, cardiomyocytes, and megakaryocytes, it is also highly prevalent in cancer, where it represents one of the most frequent large-scale genomic alterations across tumor types^4-6^. Notably, WGD is strongly enriched in *TP53*-mutant cancers, indicating selective pressure to disable p53-dependent surveillance mechanisms that would otherwise limit the proliferation of genome-doubled cells^7-9^. Indeed, for several decades WGD has been recognized as a potent trigger of p53 activation, leading to the postulation of a “tetraploidy checkpoint” that restrains the expansion of polyploid cells^10-13^. More recent work has refined this concept by demonstrating that mammalian cells do not directly sense genome content *per se* but instead rely on centrosome amplification as a structural proxy for genome doubling. Because WGD is typically accompanied by centrosome duplication, the presence of supernumerary centrosomes provides a robust and spatially localized cue that engages dedicated stress-response pathways^14,15^. This centrosome-based surveillance mechanism has been extensively validated *in vitro* and is supported by *in vivo* evidence from physiologically polyploid tissues, including hepatocytes, where centrosome number correlates with ploidy and elicits p53 activity^14-22^.

Caspase-2 has emerged as a key effector of this surveillance response. An evolutionarily conserved member of the caspase family, Caspase-2 functions predominantly in a non-apoptotic, tumor-suppressive role in this context^23^. Upon centrosome-associated stress following WGD, Caspase-2 is activated by the PIDDosome, a multiprotein complex that assembles at centrosomes and promotes Caspase-2 trans-autoproteolysis^14,24^. Activated Caspase-2 cleaves the E3 ubiquitin ligase MDM2, abrogating its ligase activity and stabilizing p53, thereby linking centrosome-based sensing of genome doubling to p53-dependent transcriptional responses^14,25^. While localization of the PIDDosome subunit PIDD1 to distal appendages of the mother centriole is required to trigger Caspase-2 activation^16,18^, whether additional centrosomal components or architectural features contribute to the spatial regulation of this cascade remains unclear.

Despite a central role of Caspase-2 in the cellular response to WGD, interrogating this pathway in living cells has remained challenging. Activation of Caspase-2 following WGD elicits dynamic transcriptional responses already during the first post-WGD cell cycle^20^, a process that cannot be readily captured by existing assays. Current approaches to monitor Caspase-2 activity rely largely on fluorogenic peptide substrates that lack specificity and are efficiently cleaved by multiple caspases, particularly in apoptotic settings, precluding discrimination of Caspase-2 activity associated with genome surveillance from broader cell death programs^26,27^. Genetically encoded proximity-based sensors, such as bimolecular fluorescence complementation (BiFC) reporters, detect Caspase-2 oligomerization but report on protein proximity rather than proteolytic activity^28-30^. Consequently, tools enabling a direct and functional readout of Caspase-2 pathway engagement in a post-WGD context remain lacking.

The dynamic regulation of RNA modifications, collectively referred to as the epitranscriptome, represents an important layer of post-transcriptional control in cellular stress responses and tumor suppression^31,32^. Among these modifications, N6-methyladenosine (m6A) is the most abundant internal mark on mRNA and is deposited by a multisubunit methyltransferase complex -commonly referred to as the m6A writer-comprising the catalytic subunit METTL3 together with associated adaptor and regulatory proteins^33^. Components of the m6A writer complex have been implicated in shaping p53-dependent transcriptional programs, particularly in response to genotoxic stress, suggesting that epitranscriptomic regulation may influence genome surveillance pathways^34-36^. However, whether m6A-dependent regulation contributes to signaling pathways that are rapidly engaged and dynamically rewired following WGD remains unclear.

Here, we present a genetically encoded reporter designed to functionally monitor Caspase-2 pathway activation in living cells by leveraging its endogenous substrate, MDM2. The reporter consists of a cleavage-sensitive fluorescent MDM2 fusion that reports Caspase-2-dependent MDM2 processing, providing a direct readout of pathway output rather than enzymatic activity in isolation. Using this reporter in combination with high-throughput small-molecule screening and genome-wide CRISPR knockout screening, we dissect regulatory layers acting upstream and downstream of Caspase-2 activation. This strategy reveals rewiring of the p53-MDM2 circuit following WGD, uncovers a state-dependent role for epitranscriptomic regulation mediated by the m6A writer complex, and identifies centrosomal architectural features that govern PIDDosome-dependent pathway activation.

## Results

### Endogenous tagging of MDM2 with mScarlet enables detection of Caspase-2 activity

To establish a cellular system suitable for monitoring Caspase-2 activity, we screened a panel of human cell lines for accumulation of the Caspase-2-dependent MDM2 cleavage product in response to WGD. Inhibition of Aurora B kinase with ZM447439 (ZM)^14,16,37^ induced detectable MDM2 cleavage in multiple lines, with human breast carcinoma cell line Cal51 showing the most robust response (Fig. S1A). RNA-seq analysis further revealed p53-dependent transcriptional changes in ZM-treated Cal51 cells, which were largely abolished in *TP53* knockout derivatives (Fig. S1B-C). Together, these data establish Cal51 as a suitable model to investigate the Caspase-2-MDM2-p53 signaling axis.

The N-terminal MDM2 fragment generated by Caspase-2 cleavage lacks E3 ligase activity and accumulates due to increased stability. Consistent with this, cycloheximide chase experiments in ZM-treated Cal51 cells showed that the cleavage fragment persists markedly longer than the full-length protein (Fig. S1D). By leveraging this property, we generated a Cal51 knock-in line expressing an N-terminal mScarlet-tagged MDM2 from the endogenous locus^38^, immediately downstream of the second translation initiation codon. The tag includes a V5 epitope, enabling detection of both full-length and cleaved MDM2 species (Fig. 1A). PCR-based genotyping confirmed successful homozygous tagging (Fig. S1E), and immunoblot analysis showed that N-terminal insertion of the V5-mScarlet tag preserves Caspase-2-dependent MDM2 cleavage and p53 accumulation (Fig. 1B). We refer to this engineered line as Cal51^*mScarlet-MDM2*^ and validated two independent clones for correct targeting and functional responsiveness. Unless otherwise indicated, experiments were performed using clone #1.

**Figure 1.**
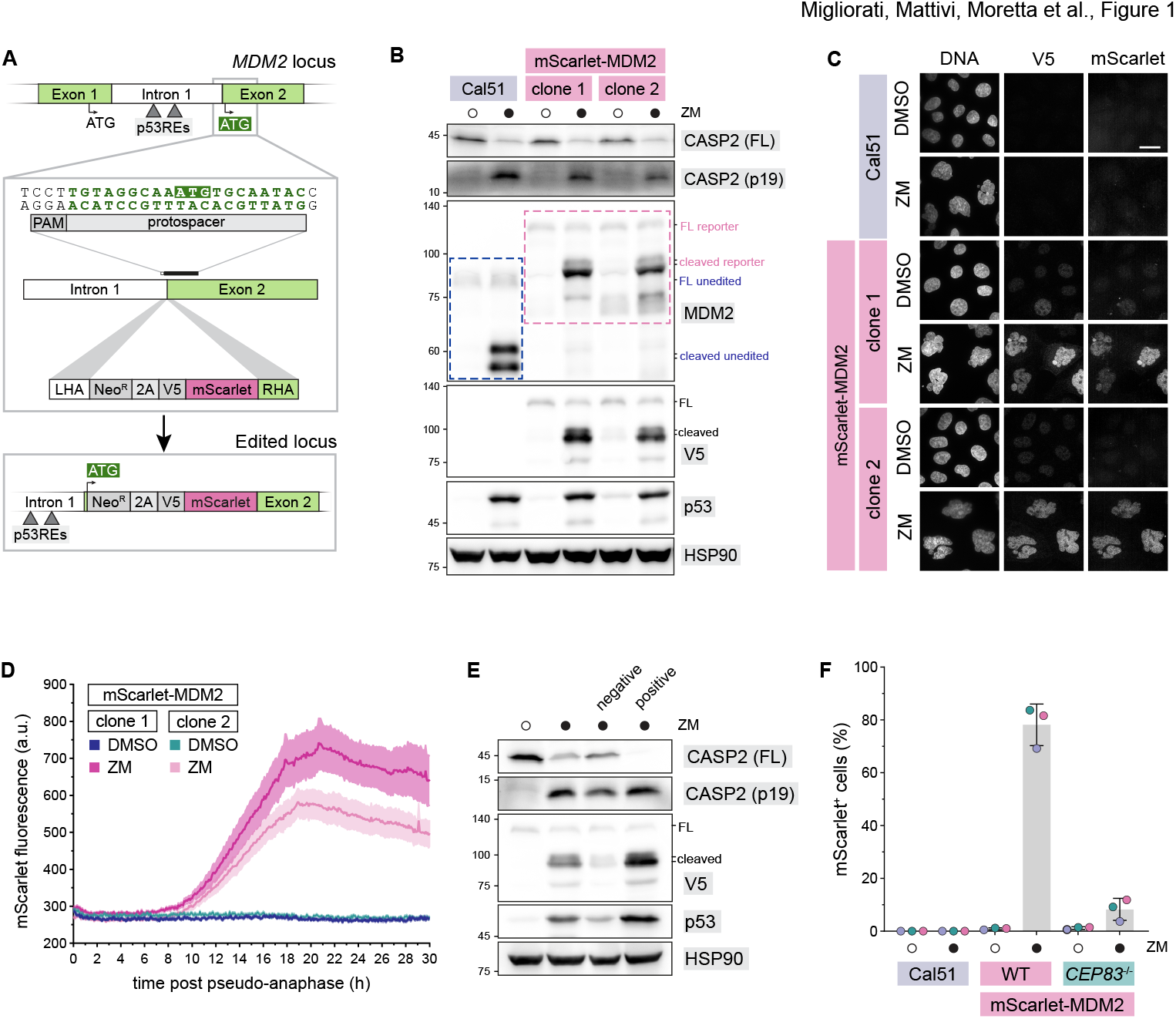
A fluorescent MDM2 reporter for Caspase-2 activation upon centrosome amplification. **(A)** Schematic representation of the V5-mScarlet knock-in at the endogenous *MDM2* locus in Cal51 cells. The tag was inserted downstream of the second translation initiation codon (ATG) and the p53-responsive elements (p53REs). The protospacer adjacent motif (PAM) and protospacer sequences are reported. LHA: left homology arm; NeoR: neomycin resistance; RHA: right homology arm. **(B)** Immunoblot analysis of the Caspase-2–MDM2–p53 pathway activation in unedited Cal51 and in two independent Cal51^*mScarlet-MDM2*^ clones treated with ZM. **(C)** Immunofluorescence analysis of Cal51^*mScarlet-MDM2*^ cells treated with ZM, showing mScarlet fluorescence and anti-V5 immunostaining. Scale bar: 20 μm. **(D)** Quantification of mScarlet fluorescence over time in individual Cal51^*mScarlet-MDM2*^ cells imaged live following ZM or vehicle-only treatment. Traces were aligned by setting time zero to the onset of pseudo-anaphase for each cell. The plot shows the population mean ± s.e.m.; n = 25 cells per condition. **(E)** Immunoblot analysis of mScarlet-positive and mScarlet-negative Cal51^*mScarlet-MDM2*^ cells isolated by FACS following ZM treatment. **(F)** Flow cytometry analysis of mScarlet fluorescence in unedited Cal51, wild-type (WT), and CEP83^-/-^ Cal51^*mScarlet-MDM2*^ cells following ZM treatment.

We next evaluated the performance of the system by assessing mScarlet fluorescence in fixed Cal51^*mScarlet-MDM2*^ cells following ZM treatment. As expected, fluorescent signal accumulated in most treated cells, consistent with Caspase-2-dependent stabilization of the cleaved MDM2 fragment (Fig. 1C). To characterize the dynamics of this response, we performed time-lapse imaging and quantified mScarlet accumulation in individual cells. In ZM-treated cells, fluorescence began to increase approximately 8 h after pseudo-anaphase onset and continued to rise gradually, reaching a maximum around 18–20 h (Fig. 1D, S1F, and Supplementary Movie 1). To confirm that mScarlet fluorescence accurately reports Caspase-2 activation, we performed fluorescence-activated cell sorting (FACS) of ZM-treated cells based on reporter signal and analyzed sorted populations by immunoblot. Fluorescent cells showed robust Caspase-2 autoproteolysis and MDM2 cleavage, whereas non-fluorescent cells did so to a much lesser extent (Fig. 1E and S1G), validating the ability of the reporter to distinguish MDM2-cleavage competent from non-competent cells within the same treatment condition. To further assess reporter specificity, we targeted CEP83, a distal appendage component required for recruitment of PIDD1 to mature mother centrioles and subsequent Caspase-2 activation^16^. CRISPR-mediated knockout of *CEP83* in Cal51^*mScarlet-MDM2*^ cells abolished MDM2 cleavage and mScarlet fluorescence upon ZM treatment (Fig. 1F and S1H), confirming that reporter activation depends on distal appendage–mediated PIDDosome signaling rather than on centrosome amplification *per se*.

Collectively, these findings demonstrate that mScarlet fluorescence accumulation in Cal51^*mScarlet-MDM2*^ cells faithfully reflects endogenous Caspase-2 activity downstream of centrosome amplification, providing a genetically encoded biosensor of pathway output at single-cell resolution.

### High-content imaging reveals differential inhibition of Caspase-2 by pan- and selective caspase inhibitors

Having validated the reporter as a faithful readout of Caspase-2 pathway activation, we next leveraged it to profile the activity of six caspase inhibitors in a high-content dose-response format. The inhibitory potency (IC_50_) of the selected compounds towards Caspase-2 was determined by administering them to Cal51^*mScarlet-MDM2*^ cells together with ZM and quantifying reporter fluorescence across a log_2_ dilution series (0.312-20 μM). In parallel, the same compounds were tested in non-engineered Cal51 cells to assess inhibition of apoptotic executioner caspases using a commercial DEVD-based fluorescent substrate, following a pro-apoptotic treatment (staurosporine + the BH3 mimetic ABT-737). This dual setup allowed us to assess both the potency and selectivity of each compound across two distinct caspase activation contexts. Consistent with the distinct activation profiles induced by ZM and STS + ABT-737 (Fig. S2A-B), we observed marked differences in the activity of the inhibitors between the two assay conditions.

Among the compounds tested, the first-in-class pan-caspase inhibitor Emricasan (also known as IDN-6556)^39^ exhibited potent, dose-dependent inhibition in both assays, emerging as the most effective caspase inhibitor in our screen (Fig. 2A-D). The poly-caspase inhibitor Q-VD-OPh (QVD)^40,41^ suppressed apoptotic caspase activation but showed minimal activity against Caspase-2, highlighting the limited efficacy of some broadly used inhibitors on this specific caspase (Fig. 2C-D and Fig. S2C-D). The peptidomimetic compounds LJ2a and LJ3a selectively inhibited Caspase-2 activity, while having negligible effects on apoptotic caspases, as previously reported^42^. LJ3b, an inactive stereoisomer of LJ3a, showed no detectable activity in either context. Belnacasan (also known as VX-765), a preferential inhibitor of human Caspase-1, -4, -5, also failed to elicit any measurable response, in line with its reported lack of efficacy toward non-inflammatory caspases^43^ (Fig. 2C-D and Fig. S2C-D).

**Figure 2.**
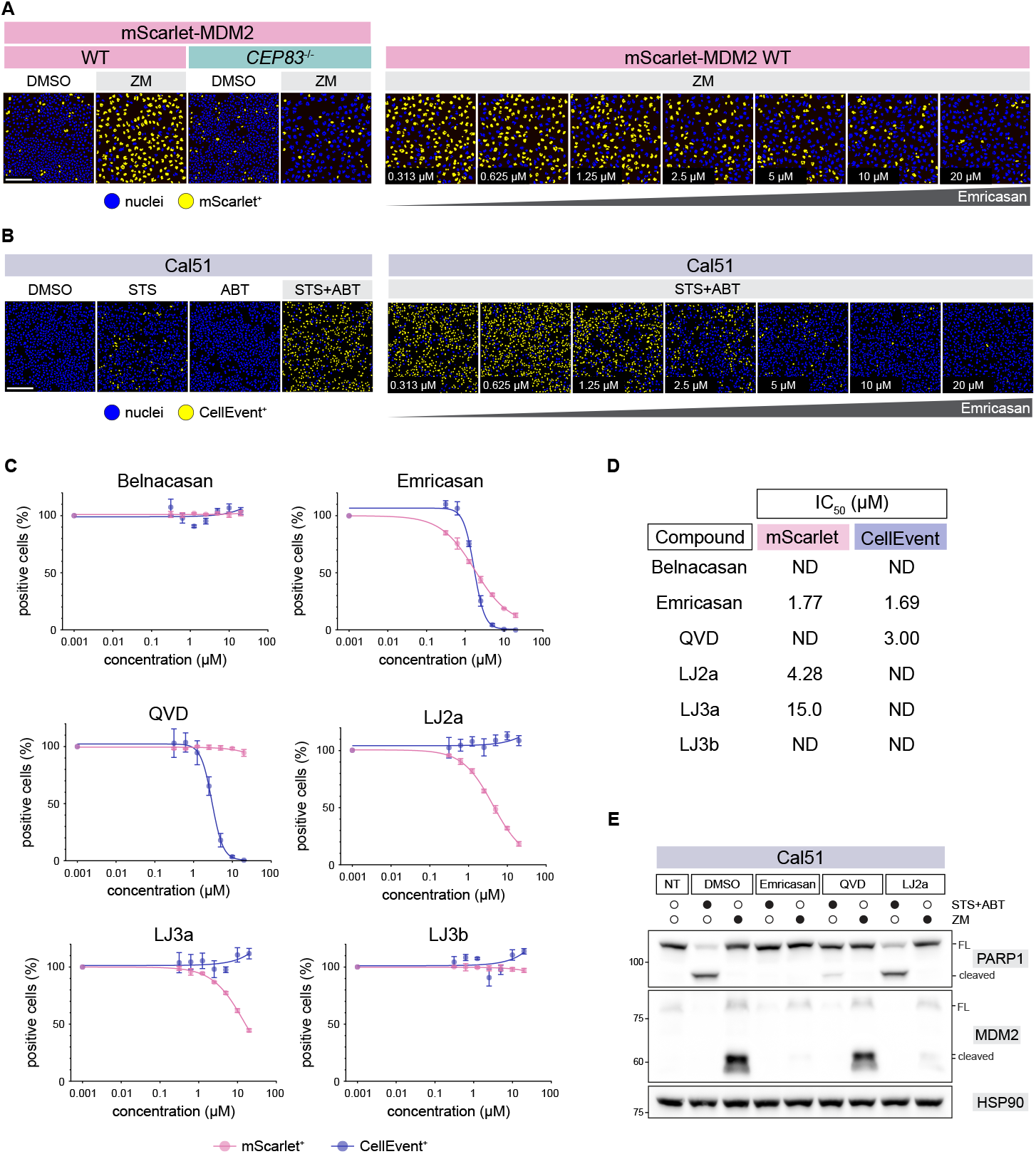
High-content screening of caspase inhibitors reveals differential effects on Caspase‐2 and effector caspases. **(A)** Representative high-content images of Cal51^*mScarlet-MDM2*^ wild-type (WT) and CEP83^-/-^ cells treated with DMSO or ZM, and of Cal51^*mScarlet-MDM2*^ WT cells treated with ZM in the presence of increasing concentrations of Emricasan. Nuclear mScarlet fluorescence is displayed as binary labeling: nuclei exceeding a predefined fluorescence threshold are pseudo-colored in yellow, whereas nuclei below threshold are shown in blue. Scale bar: 200 μm. **(B)** Representative high-content images of Cal51 cells treated with staurosporine (STS), ABT-737 (ABT), or the combined treatment (STS + ABT), in the presence of increasing concentrations of Emricasan. Effector caspase activity was assessed using the CellEvent reporter and is displayed as binary nuclear labeling, with CellEvent-positive nuclei shown in yellow and negative nuclei in blue. Scale bar: 200 μm. **(C)** Dose-response curves for six caspase inhibitors (Emricasan, QVD, LJ2a, LJ3a, LJ3b, and Belnacasan) measured using two high-content assays performed under distinct treatment conditions. Nuclear mScarlet fluorescence was quantified in Cal51^*mScarlet-MDM2*^ cells treated with ZM, whereas effector caspase activity was quantified in Cal51 cells treated with STS + ABT using the CellEvent reporter. Data represent 3 biological replicates, each with 3 technical replicates (n = 9). **(D)** Table summarizing IC_50_values for the indicated caspase inhibitors obtained from the two assay conditions shown in (C). ND, not determined. **(E)** Immunoblot analysis of Cal51 cells treated with ZM or with STS + ABT in the presence of vehicle (DMSO), Emricasan, QVD, or LJ2a (10 μM each).

To validate these findings biochemically, we treated Cal51 cells with a subset of inhibitors (Emricasan, QVD, and LJ2a) under both caspase activation conditions and analyzed lysates by immunoblotting for MDM2 and PARP1 cleavage - readouts of Caspase-2 and effector caspase activity, respectively. In agreement with the screening data, QVD selectively blocked PARP1 cleavage, LJ2a suppressed MDM2 processing, and Emricasan inhibited both Caspase-2 and executioner caspases (Fig. 2E). Finally, enzymatic inactivation parameters (*k*_*inact*_/K_I_) measured using recombinant human Caspase-2 and Caspase-3 were consistent with the differential cellular effects observed for Emricasan, QVD, and LJ2a (Fig. S2E-F).

Together, these results establish our engineered reporter system as a robust platform for compound screening and mechanism-of-action studies, enabling simultaneous assessment of caspase selectivity and potency in a physiologically grounded cellular context.

### Kinome-wide screening implicates PLK1 in Caspase-2 activation downstream of centrosome duplication

We next leveraged the reporter to systematically identify regulators of Caspase-2 activation in response to WGD through a kinome-wide high-content screen using the KCGS library of annotated kinase inhibitors^44^. Cal51^*mScarlet-MDM2*^ cells were treated with ZM in combination with each compound, and reporter fluorescence was quantified by automated microscopy. To distinguish compounds that suppress Caspase-2 signaling from those that impair cell viability, Z-scores for mScarlet-positive nuclei were plotted against total cell number. This analysis revealed a distinct subset of six compounds that selectively suppressed reporter activation without reducing cell counts (Fig. 3A and Fig. S3A). Notably, all six compounds shared annotated activity against PLK1 (Fig. 3B), implicating PLK1 as a candidate regulator of Caspase-2 activation under WGD promoting conditions.

**Figure 3.**
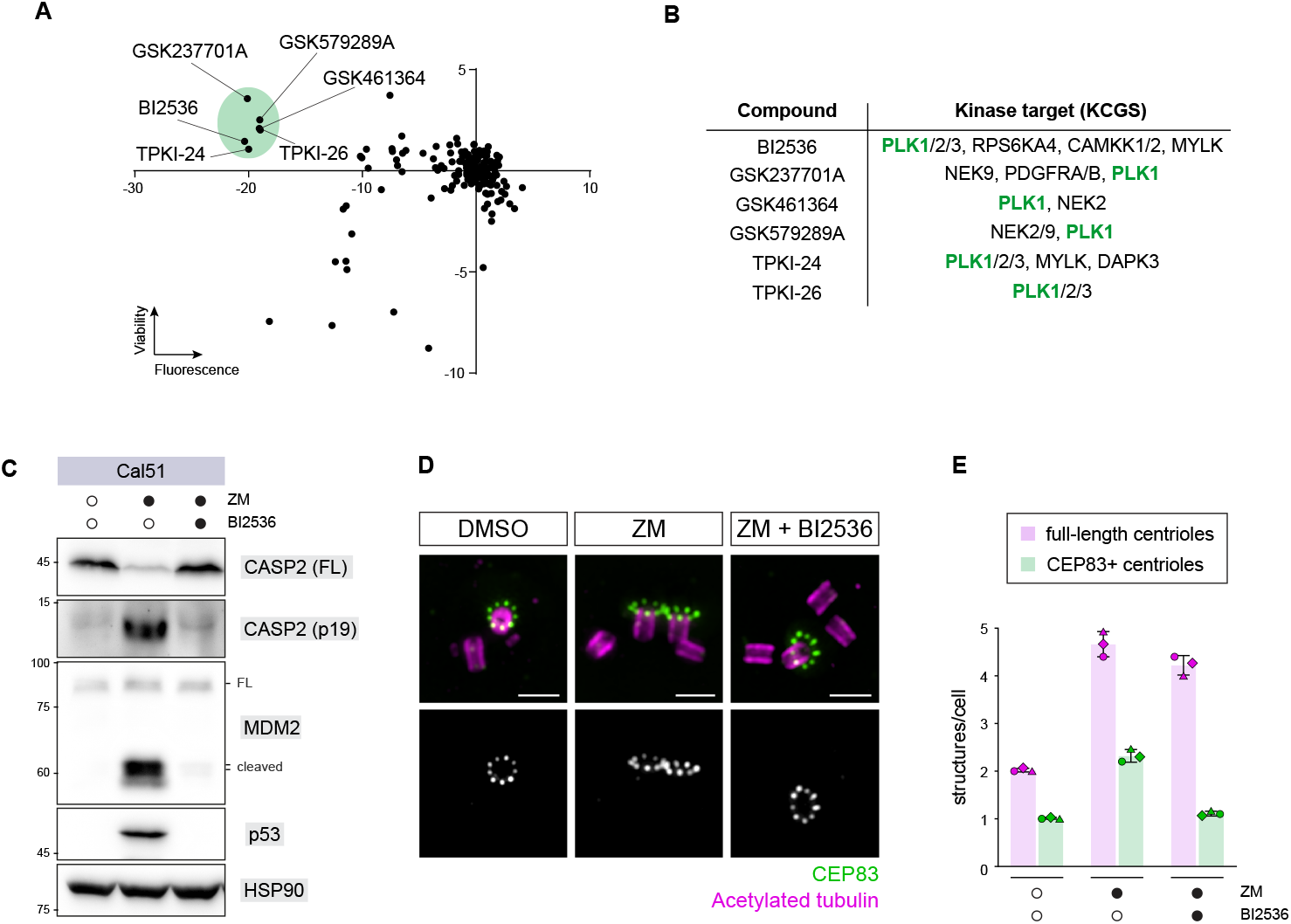
Kinome-wide screening implicates PLK1 in Caspase-2 activation by regulating centriole maturation. **(A)** High-content screen of the KCGS kinase inhibitor library performed in Cal51^*mScarlet-MDM2*^ cells co-treated with ZM. Z-scores for the fraction of nuclei exhibiting mScarlet fluorescence are plotted against Z-scores for total cell number. Compounds that reduced the fraction of mScarlet-positive nuclei without a concomitant reduction in cell number are highlighted. **(B)** Table listing the six compounds identified as hits in (A) and their annotated kinase targets based on KCGS profiling data. All six compounds share activity against PLK1. **(C)** Immunoblot analysis of Cal51 cells treated with ZM in the presence or absence of the selective PLK1 inhibitor BI2536. **(D)** Ultrastructure expansion microscopy (U-ExM) coupled with Structured Illumination Microscopy (SIM) was used to analyze centrosome architecture in Cal51 cells treated with ZM, in the presence or absence of BI2536. Mature mother centrioles were visualized by CEP83 staining. Representative images are shown. Scale bar: 500 nm. **(E)** Quantification of the number of mature mother centrioles (CEP83-positive structures) relative to the total centriole number (acetylated tubulin-positive structures) per cell in Cal51 = treated as indicated. Bars represent mean ± standard deviation (n = 30 cells per replicate; N = 3 independent experiments, 90 cells total).

We next validated this finding using complementary pharmacological and genetic approaches. Co-treatment of ZM-stimulated Cal51 cells with the selective PLK1 inhibitor BI2536^45^ abolished MDM2 cleavage (Fig. 3C). Notably, the effects of PLK1 inhibition on Caspase-2 activation were not associated with increased mitotic arrest under polyploidizing conditions (Fig. S3B). To exclude off-target effects, we leveraged a human hTERT-RPE1 (RPE1) cell line expressing an analog-sensitive allele of PLK1 (PLK1^as^)^46^. Acute inhibition of PLK1^as^ with the ATP analog 3MB-PP1 phenocopied the effect of BI2536, resulting in blunted MDM2 cleavage and confirming the on-target requirement for PLK1 activity (Fig. S3C).

PLK1 is known to regulate centriole maturation, including acquisition of distal appendages by mother centrioles^47-49^, structures that are required for PIDDosome assembly^16,18^. Whether PLK1-dependent remodeling of centrosomal architecture promotes the generation of assemblies capable of activating the PIDDosome under WGD has remained unclear. To address this, we used ultrastructure expansion microscopy (U-ExM)^50^ to examine centrosome architecture in ZM-treated Cal51 cells, in the presence or absence of PLK1 inhibition. As expected, ZM treatment alone resulted in a doubling of the number of both full-length centrioles and distal appendage-decorated (CEP83-positive) mother centrioles. By contrast, co-treatment with BI2536 preserved the increased number of full-length centrioles induced by ZM but restored the average number of distal appendage positive centrioles to approximately one per cell (Fig. 3D–E). Together, these results demonstrate that PLK1 activity enables the centriole maturation state required to generate a spatial configuration permissive for Caspase-2 activation following centrosome amplification displayed by WGD+ cells.

### A genome-wide CRISPR screen identifies modulators of the centrosome-PIDDosome-p53 pathway

To identify genes required for Caspase-2-dependent accumulation of the MDM2 cleavage product, we conducted a genome-wide CRISPR knockout screen in Cal51^*mScarlet-MDM2*^ cells. Following transduction with the Brunello sgRNA library^51^, cells were treated with ZM for 40 h and subsequently sorted by FACS based on fluorescence intensity. sgRNA abundance was quantified by next-generation sequencing in sorted mScarlet-low and mScarlet-high populations, and candidate regulators were ranked using MAGeCK^52^ (Fig. 4A).

**Figure 4.**
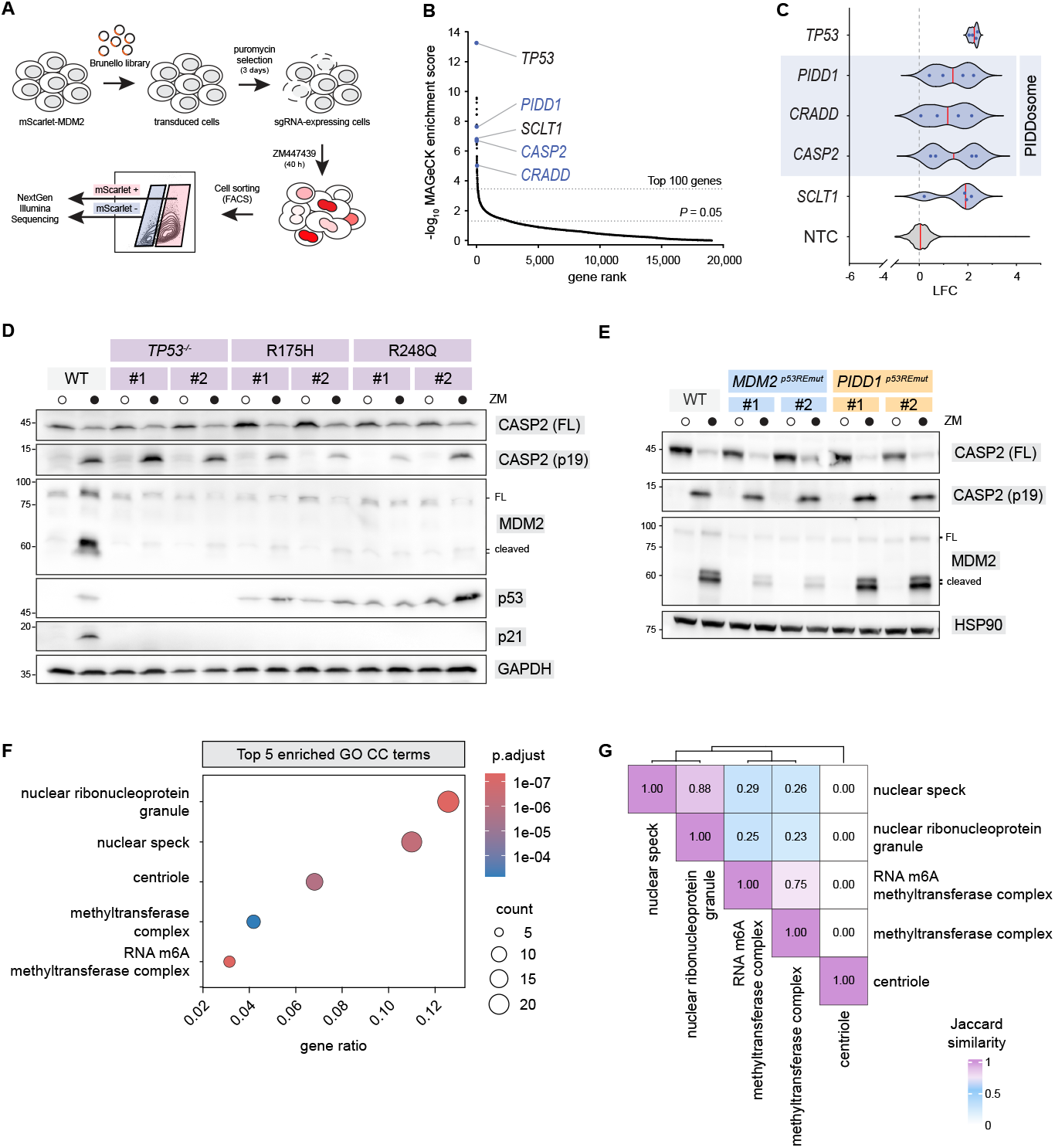
A genome-wide CRISPR screen identifies modulators of Caspase-2–dependent MDM2 cleavage. **(A)** Schematic overview of the genome-wide CRISPR knockout screen in Cal51^*mScarlet-MDM2*^ cells. Cells were transduced with the Brunello sgRNA library, subjected to puromycin selection, treated with ZM for 40 h, and sorted by FACS based on reporter fluorescence. Genomic DNA from sorted populations was processed for sgRNA amplification and Illumina-based next-generation sequencing. **(B)** Waterfall plot showing genome-wide ranking of genes based on MAGeCK analysis of sgRNA enrichment in the reporter-negative population. Genes are ordered by rank, and the y-axis represents the -log10-transformed MAGeCK enrichment score (robust rank aggregation score). Selected pathway components are highlighted. **(C)** sgRNA-level log_2_fold-change (LFC) values for selected genes identified in the CRISPR screen. Each point represents an individual sgRNA from the Brunello library targeting the indicated gene, illustrating the consistency of sgRNA behavior across hits. NTC: non-targeting control. **(D)** Immunoblot analysis of an isogenic panel of RPE1 cells comprising wild-type p53 cells (WT), *TP53*^-/-^ cells, or cells expressing hotspot mutant p53 alleles (R175H or R248Q) following ZM treatment. Two independent clones are shown for each *TP53*^-/-^, R175H, and R248Q genotype. **(E)** Immunoblot analysis of wild-type (WT) and CRISPR-engineered Cal51 cells harboring point mutations disrupting the p53 response elements (p53REs) in the *PIDD1* or *MDM2* promoters, as schematized in Fig. S4A, following ZM treatment. Two independently derived clonal cell lines are shown for each genotype. **(F)** Dot plot of Gene Ontology (GO) Cellular Component terms enriched among the top 200 genes identified in the CRISPR screen. Dot size represents the number of genes associated with each term, and color indicates enrichment significance. **(G)** Jaccard similarity heatmap showing pairwise overlap between the five top-ranked enriched GO Cellular Component terms shown in (F). Each square reports the Jaccard similarity index, and colour intensity reflects similarity magnitude, highlighting extensive overlap among m6A writer-associated terms and no overlap with the centriole term.

Consistent with the specificity of the screen, all three PIDDosome components - *PIDD1, CRADD* (RAIDD), and *CASP2*-were among the top-ranked hits, together with *SCLT1*, which encodes a distal appendage protein required for PIDD1 recruitment to the centrosome^16^ (Fig. 4B-C). Notably, *TP53* emerged as the top-ranked gene, despite being canonically positioned downstream of MDM2 cleavage. This finding raised the possibility that p53 may engage in a positive feedback loop to amplify or stabilize the Caspase-2 pathway output specifically upon WGD.

To investigate the role of p53 in more detail, we used an isogenic panel of RPE1 cells in which the endogenous *TP53* gene was either knocked out or replaced by CRISPR-mediated knock-in with common hotspot mutations (R175H or R248Q) ^53^ that cause transcriptional inactivity of *TP53*. Both loss of p53 and expression of hotspot p53 mutants abolished accumulation of the Caspase-2–generated MDM2 cleavage product upon ZM treatment, demonstrating that p53 transcriptional activity is required to sustain MDM2 cleavage (Fig. 4D).

To dissect whether this requirement reflects p53’s role in transactivating pathway components, we focused on two known transcriptional targets of p53: *PIDD1*^54^ and *MDM2*^55^. Using CRISPR-mediated editing, we selectively disrupted the p53 response elements (p53RE) in the promoter regions of each gene and validated the resulting alleles based on their transcriptional response to Nutlin-3a treatment (Fig. S4A). As expected, *PIDD1*^*p53REmut*^ and *MDM2*^*p53REmut*^ cells failed to upregulate the corresponding transcript upon p53 activation (Fig. S4B-C).

Surprisingly, despite impaired p53-dependent transactivation of *PIDD1, PIDD1*^*p53REmut*^ cells retained Caspase-2 activity and robust MDM2 cleavage. By contrast, cells with disrupted *MDM2* transcriptional activation failed to accumulate the MDM2 cleavage product, despite showing normal Caspase-2 autoproteolysis, revealing a positive feedback mechanism in which p53 sustains the expression of its own cleavage-dependent activator (Fig. 4E).

These results highlight an important interpretive feature of the CRISPR screen: because the reporter signal ultimately depends on the accumulation of cleaved MDM2, enriched hits can reflect regulators acting at different levels of the Caspase-2-MDM2-p53 axis. To organize this functional diversity, we performed a Gene Ontology (GO) over-representation analysis (ORA) focused on the Cellular Component category, using the top 200 hits from the screen as input. For clarity and downstream validation, we focused on the five top-ranked enriched terms, which clustered into two groups based on gene overlap (Fig. 4F-G). Four terms, -”nuclear ribonucleoprotein granule”, “nuclear speck”, “methyltransferase complex”, and “RNA N6-methyladenosine methyltransferase complex”-showed extensive mutual overlap, driven by the shared inclusion of m6A-writer components across these GO terms (Fig. 4G). The remaining term - “centriole”-displayed no overlap with the other ontologies, highlighting the presence of a genetically distinct centrosome-associated module. This modular separation provides a clear rationale for the two mechanistically distinct routes of functional follow-up pursued in this study.

### The m6A writer complex gates p53-MDM2 feedback circuit output

Given the coordinated enrichment of m6A writer components in the CRISPR screen, we next investigated how this machinery contributes to pathway output. All seven core and regulatory subunits (*METTL3, METTL14, WTAP, VIRMA, ZC3H13, CBLL1*, and *RBM15*) were among the top-ranked hits (Fig. 5A), and sgRNA-level log_2_fold-change values showed consistent behavior across independent guides (Fig. 5B), indicating a coherent dependency at the level of the writer complex.

**Figure 5.**
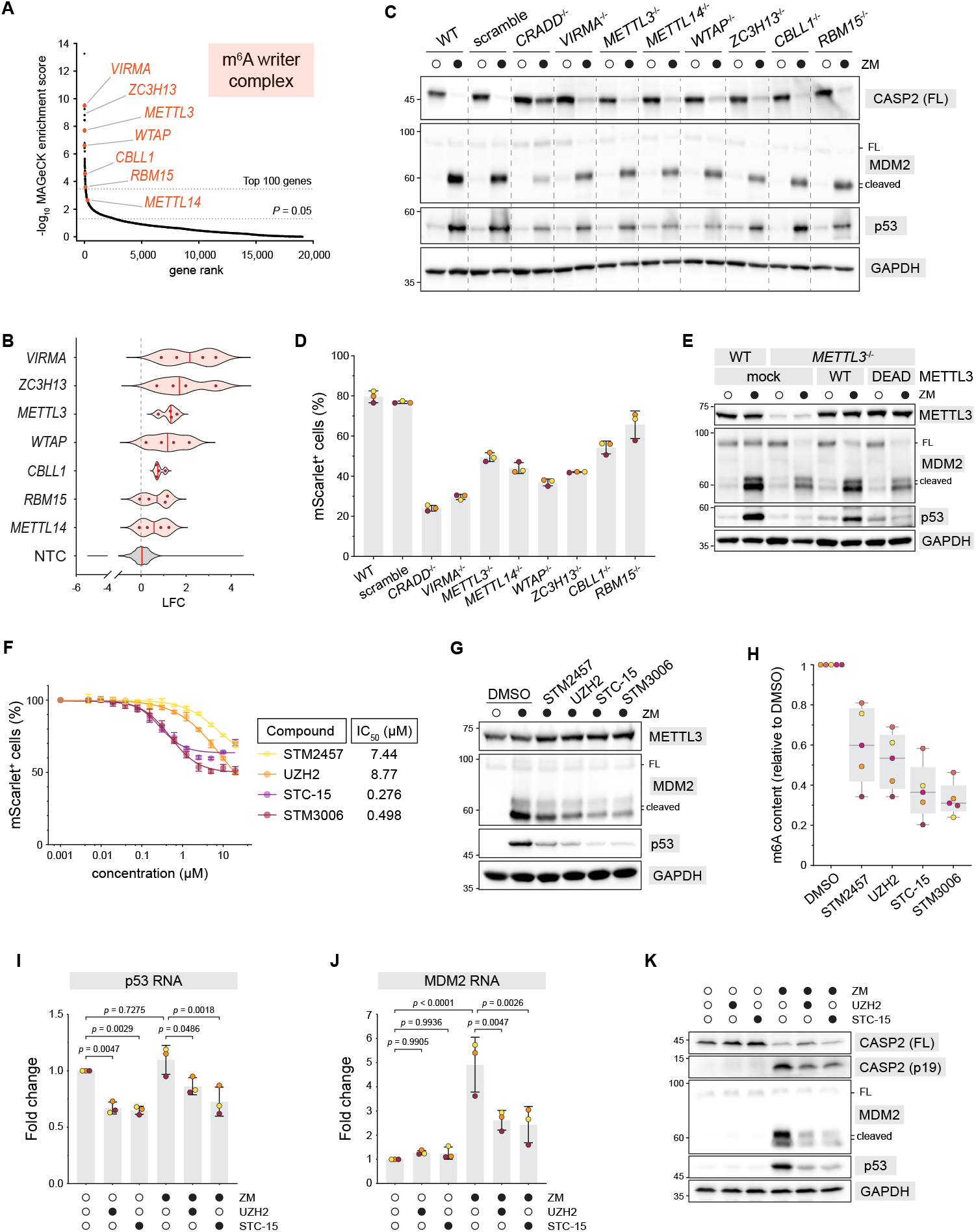
Genome-wide CRISPR screening identifies m6A writer complex components associated with modulation of the p53-MDM2 axis following WGD. **(A)** Waterfall plot of the genome-wide CRISPR knockout screen showing ranked gene enrichment based on MAGeCK analysis of sgRNA representation in the mScarlet-low population following ZM treatment. The seven core components of the m6A writer complex (*METTL3, METTL14, WTAP, VIRMA, ZC3H13, RBM15*, and *CBLL1*) are highlighted. **(B)** sgRNA-level log_2_fold-change (LFC) values for each m6A writer complex gene identified in the CRISPR screen. Each point represents an individual sgRNA from the Brunello library targeting the indicated gene. NTC: non-targeting control. **(C)** Immunoblot analysis of Cal51 cells following individual knockout of each m6A writer complex component and ZM treatment. *CRADD* (RAIDD) knockout is shown as a positive control for suppression of pathway output. **(D)** Flow cytometry analysis of mScarlet fluorescence in Cal51^*mScarlet-MDM2*^ cells following knockout of the indicated m6A writer components and ZM treatment. Bars represent mean ± standard deviation (N = 3 independent replicates). **(E)** Immunoblot analysis of METTL3-deficient cells reconstituted with wild-type (WT) or catalytic-dead METTL3 (DPPW→APPA) following ZM treatment. **(F)** Dose-response curves for four chemically distinct METTL3 inhibitors based on nuclear mScarlet fluorescence measured in Cal51^mScarlet-MDM2^ cells following ZM treatment. IC_50_values were estimated for each compound. **(G)** Immunoblot analysis of Cal51 cells treated with ZM and the indicated METTL3 inhibitors (each at 2.5 µM). **(H)** Global m6A levels measured by ELISA on poly(A)-enriched mRNA isolated from Cal51 cells following treatment with METTL3 inhibitors. Box plots display medians (horizontal lines), the interquartile range (grey boxes), maximum-to-minimum range (whiskers) and individual data points (N = 5 independent replicates); m6A content is normalized to vehicle-treated cells. **(I-J)** RT–qPCR analysis of p53 (I) and MDM2 (J) transcript levels in Cal51 cells under basal conditions and following ZM treatment, in the presence or absence of METTL3 inhibition. Transcript levels are shown relative to the untreated condition. One-way ANOVA test. **(K)** Immunoblot analysis corresponding to the samples shown in (I–J).

To functionally validate this dependency, acute disruption of any m6A writer subunit markedly reduced accumulation of the Caspase-2-generated MDM2 cleavage fragment following pathway activation, despite preserved Caspase-2 activation, and was accompanied by diminished p53 protein levels (Fig. 5C and S5A). Consistent with these bulk effects, m6A writer disruption also resulted in a pronounced loss of reporter fluorescence at the single-cell level (Fig. 5D and S5B), establishing the writer complex as a determinant of pathway output rather than Caspase-2 enzymatic activity *per se*. Notably, centrosome integrity was preserved upon m6A writer disruption, consistent with intact upstream pathway initiation (Fig. S5C–D).

To determine whether this requirement depends on the catalytic activity of the writer complex, rescue experiments in METTL3-deficient cells demonstrated that re-expression of wild-type METTL3 restored accumulation of both p53 and the MDM2 cleavage product, whereas a catalytic-dead mutant (DPPW→APPA)^56^ failed to do so (Fig. 5E and S5E).

To further assess whether this dependency is pharmacologically tractable, we next inhibited METTL3 using four chemically distinct compounds^57-60^. All suppressed reporter output with varying potency (Fig. 5F and S5F), and inhibitor potency correlated with reduced accumulation of the MDM2 cleavage product and p53 protein levels (Fig. 5G), as well as with reductions in global m6A levels measured by ELISA on poly(A)-enriched mRNA (Fig. 5H), confirming on-target inhibition.

We next examined how m6A writer inhibition affects the p53-MDM2 circuit under basal and WGD conditions. In untreated cells, inhibition of METTL3 significantly reduced p53 mRNA abundance, without altering MDM2 transcript levels (Fig. 5I-J), consistent with a primary effect on p53 transcription or stability. As expected, ZM treatment alone did not affect p53 transcript levels but induced a ∼5-fold increase in MDM2 mRNA, accompanied by Caspase-2 activation and MDM2 cleavage at the protein level (Fig. 5I-K). Notably, co-treatment with METTL3 inhibitors markedly attenuated the ZM-induced increase in MDM2 mRNA, resulting in substantially reduced accumulation of the MDM2 cleavage product and diminished p53 protein levels (Fig. 5J-K).

Together, these findings position the m6A writer complex as a catalytic and pharmacologically tractable determinant of p53–MDM2 circuit output following WGD. Rather than directly modulating Caspase-2 activation, m6A-dependent regulation sustains accumulation of the MDM2 cleavage product and reinforcement of p53 signaling once the circuit has transitioned into a self-amplifying state. In this framework, epitranscriptomic control acts as a competence layer that stabilizes pathway output downstream of centrosome-triggered Caspase-2 activation.

### Centrosomal integrity enables PIDDosome function independently of distal appendage formation

Having established the transcriptional and circuit-level branches of the screen, we returned to the centrosome-associated module to define which structural features are required for PIDDosome-dependent pathway activation. To this end, we focused on a set of high-ranking screen hits encoding centrosomal proteins with annotated roles in centriole stability, appendage formation, and centrosome organization. These included the distal appendage protein SCLT1, a known regulator of PIDDosome signaling^16^, as well as multiple proteins associated with subdistal appendages or centrosome architecture, including CEP128, ODF2, NIN (Ninein), CEP350, CEP120, and TEDC1 (Fig. 6A–B)^61-66^.

**Figure 6.**
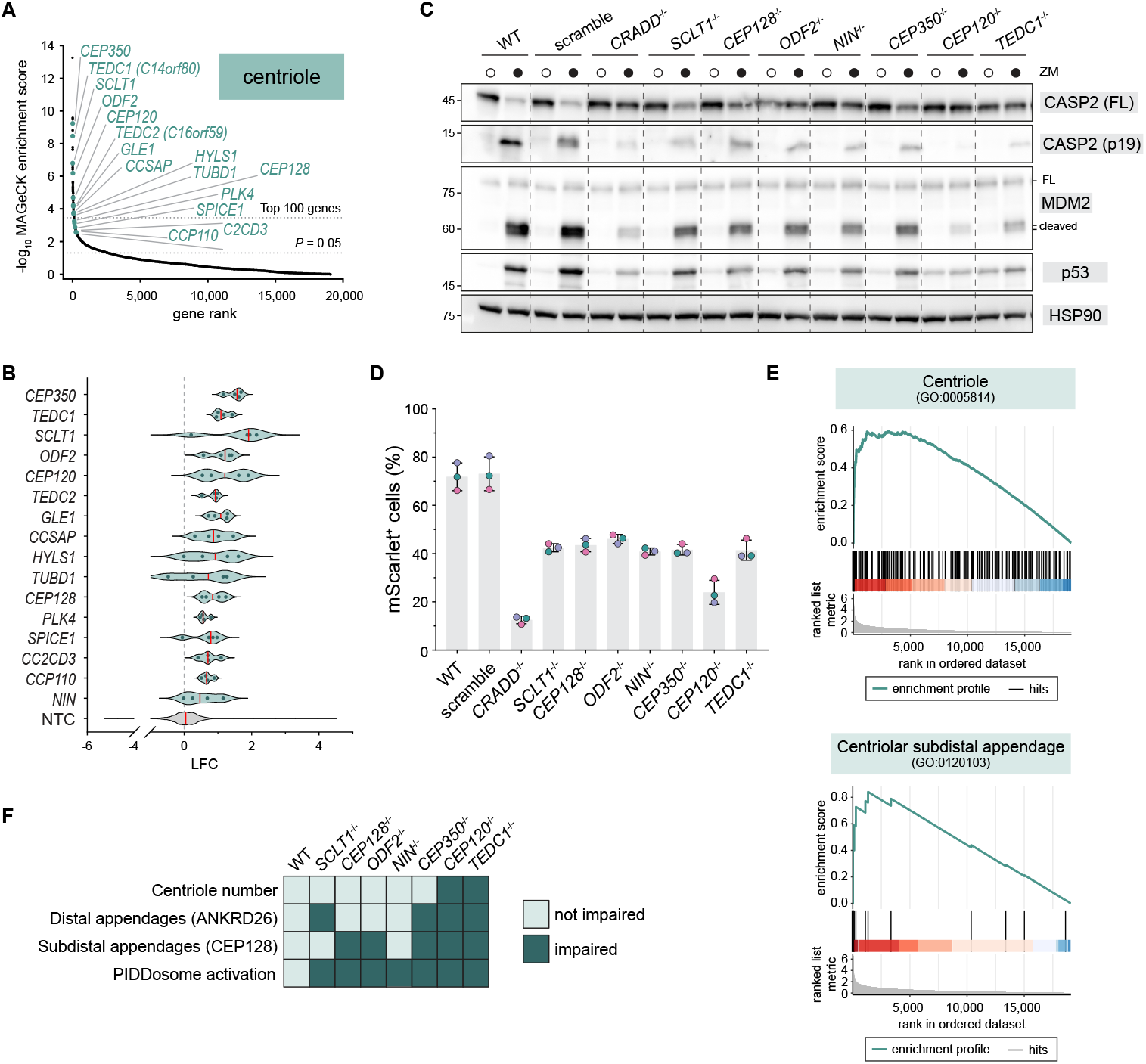
Genome-wide CRISPR screening identifies centrosome-associated genes whose disruption alters pathway output following WGD. **(A)** Waterfall plot derived from the genome-wide CRISPR knockout screen performed in Cal51^*mScarlet-MDM2*^ cells, showing ranked gene enrichment based on MAGeCK analysis of sgRNA representation in the mScarlet-low population following ZM treatment. High-ranking centrosome-associated genes, including *SCLT1, CEP128, ODF2, NIN, CEP350, CEP120*, and *TEDC1*, are highlighted. **(B)** sgRNA-level log_2_fold-change (LFC) values for the indicated centrosome-associated genes identified in the CRISPR screen shown in (A). Each point represents an individual sgRNA from the Brunello library targeting the indicated gene. NTC: non-targeting control. **(C)** Immunoblot analysis of Cal51 cells following acute disruption of the indicated centrosome-associated genes and ZM treatment. *CRADD* (RAIDD) knockout is included as a positive control for suppression of pathway output. **(D)** Flow cytometry analysis of mScarlet fluorescence in Cal51^*mScarlet-MDM2*^ cells following knockout of the indicated centrosome-associated genes and ZM treatment. The bars indicate the mean ± standard deviation (N = 3 independent replicates). **(E)** Rank-based gene set enrichment analysis of the genome-wide CRISPR screen identifying centrosome-related Gene Ontology Cellular Component terms. Centriole-associated categories are broadly represented, with subdistal appendage–related terms exhibiting high enrichment score. **(F)** Summary of centrosome-associated structural features and PIDDosome pathway proficiency following disruption of the indicated genes. The heatmap categorizes phenotypes as impaired (dark green) or not impaired (light green) for centrosome number, distal appendages and subdistal appendages integrity, and PIDDosome activation. For subdistal appendage integrity, CEP128 was used as a structural marker; accordingly, disruption of NIN (Ninein), a subdistal appendage component acting epistatically downstream of CEP128, does not score as impaired in this structural assessment^63^. Based on data displayed in panels 6C-D and S6C, integrating biochemical, flow cytometry, and visual scoring readouts.

Consistent with their enrichment in the CRISPR screen, acute disruption of each of these factors resulted in a marked reduction of pathway output, as assessed by loss of the Caspase-2-generated MDM2 cleavage product and decreased reporter fluorescence (Fig. 6C–D). In contrast to acute disruption of the m6A writer complex, these perturbations also impaired Caspase-2 activation itself, indicating that centrosomal components act at the level of pathway initiation rather than output amplification. Notably, pathway inhibition was observed across genes with diverse centrosomal functions, raising the question of which centrosomal features are required for pathway initiation.

To identify shared structural features underlying this phenotype, we performed rank-based gene set enrichment analysis across the full CRISPR dataset. This analysis revealed significant enrichment of centrosome-related Cellular Component terms, with “centriole” emerging as a broadly enriched category. Notably, within this broader group, “centriolar subdistal appendage” showed a pronounced enrichment signal (Fig. 6E), pointing to a specific centrosomal submodule warranting focused structural analysis.

To further resolve the structural basis of pathway inhibition at the level of centrosome organization, we examined the impact of these perturbations on centrosome number and appendage organization using established molecular markers. Loss of SCLT1 selectively impaired distal appendage integrity, whereas disruption of the subdistal appendage components -ODF, CEP128 and NIN-did not produce detectable defects in distal appendages, as assessed by ANKRD26 localization. By contrast, TEDC1 and CEP120 perturbation produced heterogeneous defects affecting both centriole number and appendage organization. In comparison, CEP350 disruption predominantly impacted appendage organization while preserving centriole number, underscoring distinct classes of centrosomal perturbations (Fig. S6A–C).

Notably, despite these divergent structural outcomes, disruption of each of these genes uniformly impaired pathway activation. In particular, pathway inhibition was observed even in conditions where centrosomes and distal appendages were preserved, indicating that neither centrosome presence nor distal appendage assembly alone is sufficient to support pathway initiation (Fig. 6F).

Together, these results define a centrosome-intrinsic architectural requirement for PIDDosome-dependent pathway initiation that cannot be explained solely by centrosome number or by the mere presence of distal appendage components. These observations prompted us to examine, at ultrastructural resolution, how centrosomal architecture influences PIDDosome recruitment and pathway activation in a canonical centrosome model.

### Subdistal appendages shape centrosome clustering to support Caspase-2 activation

Having found that multiple subdistal appendage-associated hits attenuate pathway initiation in Cal51 cells, we next switched to RPE1 cells to leverage clonal knockouts and ultrastructural readouts in a canonical centrosome model. In this system, clonal disruption of the subdistal appendage component *CEP128* strongly blunted Caspase-2 pathway activation following ZM-induced WGD, as reflected by reduced Caspase-2 processing, diminished accumulation of the MDM2 cleavage product and p53 (Fig. 7A).

**Figure 7.**
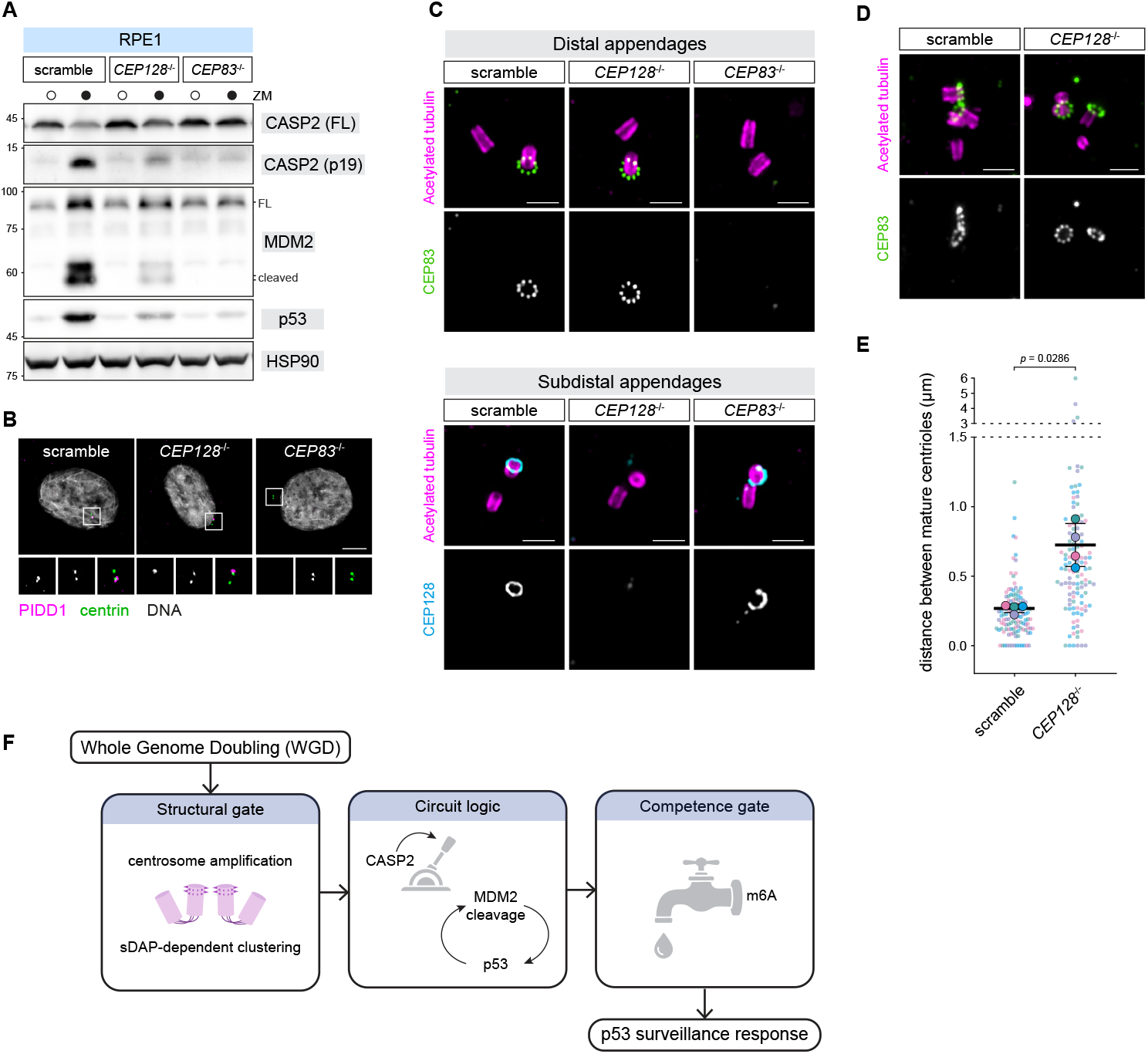
Subdistal appendage integrity impacts on centrosome clustering and pathway output following WGD. **(A)** Immunoblot analysis of RPE1 cells of the indicated genotypes following ZM treatment. **(B)** Immunofluorescence analysis of PIDD1 localization in RPE1 cells of the indicated genotypes following ZM treatment. Magnified insets of boxed regions (without the DNA channel) are shown. Scale bar: 5 μm. **(C)** Ultrastructure expansion microscopy (U-ExM) coupled with Structured Illumination Microscopy (SIM) was used to analyze distal (CEP83-positive, top panels) and subdistal (CEP128-positive, bottom panels) appendage organization in RPE1 cells of the indicated genotypes. Representative images are shown. Scale bar: 500 nm. **(D)** U-ExM/SIM analysis of centrosome spatial organization in ZM-treated RPE1 cells. Mature mother centrioles were visualized by CEP83 staining. Representative images illustrate tightly clustered or spatially separated CEP83-positive centrioles within centrosome assemblies. Scale bar: 500 nm. **(E)** Quantification of centrosome clustering in ZM-treated RPE1 cells of the indicated genotypes. The three-dimensional distance between the two closest CEP83-positive distal appendage signals was measured. Each small dot represents a single cell measurement; same-symbol colors distinguish biological replicates. The mean of each biological replicate is reported (larger dots) ± standard deviation. N = 4 biological replicates, n = 30 cells per condition; 120 cells total. Mann-Whitney test. **(F)** Proposed model for multi-layered regulation of p53 surveillance following whole-genome doubling (WGD). WGD is associated with the presence of supernumerary centrosomes that engage a centrosome-based surveillance pathway. At an upstream structural layer, subdistal appendage-dependent centrosome architecture is associated with a spatial organization of centrosomes permissive for Caspase-2 activation. Caspase-2-mediated cleavage of MDM2 alters the p53-MDM2 circuit, converting negative feedback into positive feedback and amplifying p53 signaling. At a downstream competence layer, the m6A writer complex modulates the ability of this circuit to sustain pathway output independently of Caspase-2 activation.

We then asked whether this defect reflects impaired centrosomal recruitment of the PIDDosome component PIDD1. Consistent with prior work, knockout of the distal appendage component *CEP83* led to loss of centrosomal PIDD1 localization^16^. By contrast, in *CEP128* knockout cells, PIDD1 remained robustly centrosome-associated (Fig. 7B and Fig. S7A). Thus, subdistal appendage integrity is required for efficient pathway initiation even when PIDD1 is correctly positioned at the centrosome, providing direct evidence that centrosomal PIDD1 recruitment is not sufficient to trigger Caspase-2 activation following genome doubling, even in the presence of centrosome amplification.

To explore a potential architectural basis for this requirement, we used U-ExM to assess appendage organization and centrosome spatial arrangement in RPE1 cells. Analysis of CEP83 (distal appendages) and CEP128 (subdistal appendages) organization confirmed the expected hierarchical relationship between these structures. Consistent with previous reports^67^, disruption of distal appendages altered subdistal appendage organization. While we do not resolve the distal displacement of subdistal appendages described previously, CEP83 loss was associated with a frequent transition from complete to partially organized CEP128 ring structures (Fig. 7C and Fig. S7B). By contrast, loss of subdistal appendage components did not produce detectable ultrastructural defects in distal appendage architecture. (Fig. 7C and Fig. S7C). These observations indicate that the impaired signaling observed upon subdistal appendage disruption cannot be attributed to secondary alterations in distal appendage architecture.

We next examined centrosome organization under WGD-promoting conditions. In ZM-treated control cells, supernumerary mature mother centrioles frequently assembled into tightly apposed clusters. Three-dimensional distance measurements revealed that *CEP128* knockout significantly increased inter-mother centriolar spacing, resulting in more dispersed centrosome configurations (Fig. 7D-E). This shift from compact to loosened clustering was reproducible across independent experiments and coincided with a marked loss of pathway initiation measured biochemically. Together, these data position subdistal appendages upstream of Caspase-2 activation and indicate that they promote the formation of tightly clustered centrosome assemblies permissive for PIDDosome-dependent signaling following WGD (Fig. 7F).

## Discussion

Whole-genome doubling (WGD) poses a unique challenge to cellular homeostasis, as it generates a state that is not immediately lethal, yet requires active control to prevent genotoxic stress and cellular transformation. Here, we show that surveillance of this state is governed by a multi-layered logic integrating centrosome cluster architecture, p53-signaling circuit rewiring, and epitranscriptomic competence. As such, rather than relying on a single molecular trigger, cells deploy multiple, functionally distinct layers of control that collectively determine whether p53 surveillance is initiated, amplified, and sustained following genome doubling.

Prior work established that centrosome amplification engages p53 through the PIDDosome-Caspase-2-MDM2 axis^14^ and defined distal appendages as the structural platform required for PIDDosome assembly^16,18^. These studies identified ANKRD26 as the centriolar receptor for PIDD1 and demonstrated that intact distal appendages are necessary for signaling. Importantly, ANKRD26 recruits the full-length PIDD1 precursor, whereas PIDDosome activation requires its autoproteolysis and generation of the C-terminal fragment (PIDD1-CC) that engages RAIDD^16,68^. On this basis, it was proposed that spatial proximity between mature mother centrioles might increase local concentration of PIDD1-CC and facilitate PIDDosome activation. While these findings defined the minimal structural requirements for pathway initiation, they did not address how centrosome architecture beyond distal appendage integrity contributes to signaling competence.

In this context, our identification of PLK1 as an upstream regulator provides a mechanistic link between centriole maturation and signaling competence. PLK1 activity is required to generate multiple mature mother centrioles under WGD conditions, creating a centrosomal landscape that differs qualitatively from physiological centriole configurations. Rather than merely increasing centriole number, PLK1-dependent maturation generates multiple competent mother centrioles that serve as the substrate for higher-order architectural gating.

Downstream of centriole maturation, our data identify subdistal appendage integrity as an additional requirement for efficient pathway activation. Even in the presence of properly assembled distal appendages and correctly recruited PIDD1, disruption of subdistal appendages markedly impairs PIDDosome signaling, indicating that PIDD1 localization alone is not sufficient to trigger Caspase-2 activation. Subdistal appendages have been classically characterized as microtubule-anchoring structures that contribute to centrosome positioning and ciliogenesis^63,69^, and have not previously been linked to stress sensing or p53 activation. Our findings instead indicate that signaling competence is governed by higher-order centrosomal organization, particularly the degree of clustering between mature mother centrioles. By promoting tight centrosome clustering, subdistal appendages may reinforce the spatial conditions previously proposed to facilitate local PIDD1-CC accumulation and productive PIDDosome activation.

In proliferating cells, p53 dynamics are shaped by a canonical negative feedback loop in which p53 transcriptionally induces MDM2, and MDM2 in turn promotes p53 degradation^55,70-72^. This circuit architecture can generate qualitatively distinct response modes, including oscillatory versus sustained p53 accumulation, depending on the nature and persistence of upstream inputs. Work from Lahav and colleagues established the link between feedback topology and p53 dynamics^73,74^, and further showed that Caspase-2-mediated inhibition of MDM2 can contribute to sustained p53 responses in genotoxic contexts where mitotic errors trigger PIDDosome activation^75^.

Our findings extend this model by placing PIDDosome-dependent MDM2 cleavage in a distinct structural regime. Following WGD, centrosome-triggered Caspase-2 activation converts MDM2 from a negative regulator into a cleaved substrate, thereby shifting the p53-MDM2 interaction toward a self-reinforcing state that can persist independently of transient genotoxic signals. In this context, centrosome architecture does not merely initiate a checkpoint-like response, but establishes a sustained surveillance state linked to the presence of supernumerary centrosomes. This configuration is also poised to integrate additional inputs into the p53 network, including those coming from DNA damage response (DDR) kinases, suggesting that WGD-associated structural cues and genotoxic stress pathways converge on the p53-MDM2 regulatory module to shape the qualitative nature of the p53 response.

This circuit logic further offers a basis to interpret cancer genomics observations linking WGD to selective pressures on the p53 axis. *TP53* alterations are strongly enriched in tumors that underwent WGD, whereas *MDM2* amplification-an alternative mechanism to attenuate p53 signaling in many contexts- is less prominently associated with WGD^7^. *TP53* inactivation may therefore provide a more permissive cellular state for genome-doubled cells than amplification of *MDM2*, which in the WGD context functions as a Caspase-2 substrate capable of reinforcing rather than suppressing p53 accumulation.

Beyond structural and circuit-level constraints, regulation at the level of transcript competence may further influence how genome-doubled cells sustain p53-MDM2 circuit behavior. A recent study demonstrated that METTL3 can associate with p53 protein and promote expression of selected p53 target genes, including MDM2, particularly under genotoxic stress conditions^35^. Related work further showed that RBM15 occupies the MDM2 promoter in a p53-dependent manner and supports MDM2 expression under basal conditions, indicating that components of the m6A machinery intersect with the p53–MDM2 loop beyond acute DNA damage responses^36^. In addition, the nuclear m6A reader YTHDC1 has been implicated in regulating *TP53* transcription, through mechanisms that may be partially independent of m6A deposition^34^. Together, these studies indicate that the m6A machinery interfaces with multiple nodes of the p53 network.

Whether these regulatory inputs influence the distinct structural regime imposed by WGD has remained unclear. Our data indicate that m6A writer activity regulates the p53–MDM2 circuit at multiple levels. Under basal conditions, loss of writer function reduces p53 transcript abundance, thereby influencing the availability of the circuit’s central node prior to feedback engagement. Following WGD, when Caspase-2-mediated MDM2 cleavage shifts the circuit into a self-reinforcing configuration, this upstream effect becomes functionally decisive. In this context, disruption of the writer decouples Caspase-2 activation from accumulation of the MDM2 cleavage product and selectively dampens reinforcement of p53 signaling.

In prior work, use of first-generation METTL3 inhibitors did not significantly alter *TP53* transcription at low concentrations^34^, a finding consistent with our observation that measurable effects on p53 transcript abundance require robust catalytic inhibition. Our findings do not directly address m6A deposition on specific transcripts in the WGD setting; rather, they identify a competence layer that conditions the circuit’s ability to sustain output once engaged. In this context, epitranscriptomic regulation does not simply amplify p53 target gene expression but modulates the stability and behavior of a rewired feedback topology imposed by centrosome-driven signaling.

The physiological relevance of this multilayered architecture is supported by in vivo evidence from polyploid tissues. In the murine liver, genetic ablation of *Ankrd26*, PIDDosome components, or *Tp53*, as well as expression of a non-cleavable *Mdm2* allele or loss of *Cdkn1a* (p21), all result in hyperpolyploid hepatocytes, underscoring the functional coherence of this axis in limiting genome amplification^14,17,21,22^. Notably, hepatocyte-specific deletion of *Mettl3* similarly leads to enlarged, hyperpolyploid nuclei, consistent with a requirement for m6A-dependent competence in sustaining surveillance output *in vivo*^76^.

Collectively, our findings delineate a structural-to-circuit logic in which centrosome architecture governs p53 surveillance following WGD. By linking centriole maturation and higher-order centrosomal organization to feedback rewiring of the p53-MDM2 axis, and identifying m6A-dependent regulation as a competence layer for sustained output, these findings outline how organelle structure, circuit topology, and epitranscriptomic control converge to limit genome amplification.

## Materials and methods

### Cell Culture

Cal51 (gift from Dr. Yossi Shiloh, Tel Aviv University), A549 (ATCC® CCL-185), U2OS (gift from Dr. Anna Cereseto, University of Trento), HCT116 (gift from Dr. Alessandra Bisio, University of Trento), MCF7 (DSMZ ACC 115), HepG2 (gift from Dr. Graziano Lolli, University of Trento), HEK 293T (gift from Dr. Ulrich Maurer, University of Freiburg) cell lines were cultured in DMEM (Gibco™, 11960-044). hTERT-RPE1 cells (gift from Dr. Stephan Geley, Medical University of Innsbruck) and hTERT-RPE1 PLK1^as^ cells (gift from Dr. Prasad Jallepalli, Memorial Sloan Kettering Cancer Center) were grown in DMEM/F12 1:1 (Gibco™, 11320-074). Nalm6 cells (DSMZ ACC 128) were cultured in RPMI 1640 medium (Gibco™, 31870-025). All culture media were supplemented with 10% fetal bovine serum (Gibco™, 10270-106), 2 mM L-glutamine (Gibco™, 25030-024), and a solution containing 100 IU/mL penicillin and 100 μg/mL streptomycin (Gibco™, 15140-122). Cells were maintained at 37°C in a humidified atmosphere with 5% CO_2_and were routinely screened for mycoplasma contamination.

### Drug treatments

The following compounds were used: 2 µM ZM-447439 (MCE®, HY-10128), 1 µM Staurosporine (MCE®, HY-15141), 1 µM ABT-737 (MCE®, HY-50907), 100 nM BI2536 (MCE®, HY-50698), Nutlin-3a (MCE®, HY-10029), 20 µg/Ml Cycloheximide (Thermo Scientific Chemicals, 357420010), 10 µM 3MB-PP1 (Cayman Chemical, 17860), Belnacasan (MCE®, HY-13205), Emricasan (MCE®, HY-10396), Q-VD-OPh (MCE®, HY-12305), 2.5 µM STM2457 (Cayman Chemical, 34280), 2.5 µM (Fig. 5G, 5H) or 10 µM (Fig. 5I-K) UZH2 (TargetMol®, T40357), 2.5 µM STC-15 (MCE®, HY-156677), 2.5 µM STM3006 (MCE®, HY-156773), 1 µM NU7441 (Selleckchem, S2638), LJ2a, LJ3a and LJ3b^42^ (gift from Dr. Etienne Jacotot, Paris Cité University, Inserm). Cycloheximide was dissolved in water: all the other drugs were dissolved in DMSO (Sigma-Aldrich, 472301). For ZM-447439, treatment timings were 24 h (Fig. 1B, 1C, 1E, S1A, S1B, S1D, S1G, 2E, S2B, 3C, 3D, S3B, S3C, 4D, 4E, 5C, 5G, 5I, 5J, 5K, S5A, 6C, 7A, 7D), 40 h (Fig. 1F, S1H, 4A, 5D, 5E, S5B, S5E, 6D) or 48 h (Fig. S1A). For BI2536 (Fig. 3C, 3D, S3B), 3-MB-PP1 (Fig. S3C), Nutlin-3a (Fig. S4B, S4C), STM2457 (Fig. 5G, 5H), UZH2 (Fig. 5G-K), STC-15 (Fig. 5G-K) and STM3006 (Fig. 5G, 5H), 24 h treatments were performed. Staurosporine and ABT-737 (Fig. 2E and S2B) were administered for 4 h. All untreated controls received solvent only.

### Lentivirus generation and transduction

LentiCRISPR vectors were produced as previously described^16^. Briefly, sgRNA sequences (reported in Supplementary Table S1) were cloned into the Lenti‐ CRISPR‐V2 backbone (gift from Feng Zhang; Addgene plasmid #52961). HEK 293T cells were co-transfected with the appropriate transfer plasmid together with pCMV-VSV-G (gift from Bob Weinberg, Addgene plasmid #8454) and psPAX2 (gift from Didier Trono, Addgene plasmid #12260). LentiCRISPR vectors were produced using the calcium phosphate method. Viral supernatants were collected 48 h post-transfection, filtered, and the viral titer was quantified as described previously^77^. For transduction, viral stocks were diluted in growth medium to a final concentration of 0.4 U/mL reverse transcriptase, supplemented with 4 μg/mL hexadimethrine bromide (Sigma-Aldrich, H9268), and applied to cells for 24 h. For LentiCRISPR-mediated knockout experiments targeting *CRADD, VIRMA, METTL3, METTL14, WTAP, ZC3H13, CBLL1, RBM15, SCLT1, CEP83, CEP128, ODF2, NIN, CEP350, CEP120*, and *TEDC1*, cells were selected with puromycin (2 μg/mL for Cal51; 20 μg/mL for RPE1; Invivogen, ant-pr) for 3 days and assayed 8 days post-infection. Primers used for cloning sgRNAs into the LentiCRISPR plasmid are available in Supplementary Table S1. For the generation of CL23 and CL24, CL01 and CL09 were respectively transduced with a lentiviral vector expressing the doxycycline-inducible transactivator (pLenti-ubc-rtTR-M3) and selected with 10 µg/mL blasticidin (Invivogen, ant-bl).

### CRISPR-mediated engineering of endogenous loci

The endogenous *MDM2, TP53, PIDD1*, and *CEP83* loci were engineered by nucleofection using the Lonza 4D-Nucleofector® System (Lonza Bioscience, Basel, Switzerland) following the manufacturer’s instructions and as previously described^78^. Targeting gRNAs were generated by annealing 100 μM Alt-R® CRISPR-Cas9 crRNA (Integrated DNA Technologies, IDT) with 100 μM Alt-R® CRISPR-Cas9 tracrRNA (IDT, Cat# 1072532). Ribonucleoparticles (RNPs) were assembled by mixing 150 pmol gRNA with 120 pmol recombinant Cas9 protein. After trypsinization, 2.5 × 10^5^ cells/reaction were resuspended in SE Primary Cell Full Electroporation Buffer (Lonza, Kit V4XC-1032) for Cal51 cells or P3 Primary Cell Full Electroporation Buffer (Lonza, Kit V4XP-3032) for RPE1 cells, and combined with the RNP mixture and 4 μM Alt-R® Cas9 Electroporation Enhancer (IDT, Cat# 1075915) (CL11, CL12, CL13, CL14).To generate CL09 and CL10, 200 ng of the PCR-amplified donor template were added to the electroporation mix. For all other CRISPR knock-in lines (CL15, CL16, CL17, CL18, CL19, CL20, CL21, CL22), 4 μM of the corresponding single-stranded DNA homology template (Ultramer DNA Oligonucleotide, IDT) were included. Electroporations were performed using program CH-125 (Cal51) or EA-104 (RPE1). Knockout-edited cells were recovered directly in complete medium, whereas knock-in cells were recovered in the presence of 1 μM NU-7441 for 48 h. Single cells were then seeded into 96-well plates to establish clonal lines. For CL09 and CL10 (Cal51^*mScarlet-MDM2*^), selection was performed with 800 μg/mL G418 (Invivogen, ant-gn) prior to cloning. CL12 cells were selected with 10 μM Nutlin-3a for 7 days and the resulting bulk population was used for downstream experiments. To validate editing, genomic DNA was extracted using the NucleoSpin Tissue kit (Macherey-Nagel 740952). PCR amplicons spanning the targeted loci were Sanger sequenced and analyzed with the Inference of CRISPR Edits (ICE) online tool^79^ to determine allele homozygosity. During the screening of CL15 and CL16, clones carrying frameshift indels instead of the desired base substitutions were identified and characterized as CL13 and CL14. Sequences of Alt-R® CRISPR-Cas9 crRNAs and single-stranded DNA homology template (Ultramer DNA Oligonucleotides, IDT) are available in Supplementary Table S1.

### METTL3 rescue experiment

A plasmid encoding human CRISPR/Cas9-resistant METTL3 cDNA with catalytically inactivating mutations (D395A+W398A; “APPA”) was purchased from Thermo Fisher Scientific. Reversion to the enzymatically active METTL3 sequence was achieved using Single-Primer Reactions in Parallel (SPRINP) as previously described^80^. Primer sequences used for mutagenesis are listed in Supplementary Table S1. METTL3 WT and APPA cDNAs were subcloned into FUW-tetO-MCS+ (modified from Addgene plasmid FUW-tetO-MCS #84008), and lentiviral particles were produced.

CL23 and CL24 cells were co-transduced with lentiCRISPR-METTL3 or lentiCRISPR-scramble (0.2 U/mL reverse transcriptase), together with equal titers of lentiviral vectors encoding METTL3 WT, METTL3 APPA or empty FUW-tetO-MCS+. Puromycin selection was initiated 24 h post-transduction and maintained for 3 days. Doxycycline (200 ng/mL; Thermo Fisher Scientific, 446060000) was added since the time of transduction to induce METTL3 expression. Six days post-transduction, cells were treated with 2 µM ZM-447439 for 40 h and analyzed by flow cytometry or immunoblotting.

### Immunoblotting

Cells were harvested by trypsinization, washed in PBS, lysed in lysis buffer (50 mM Tris-HCl pH 7.4, 150 mM NaCl, 0.5% NP-40, 50 mM NaF, 1 mM Na_3_VO_4_, 1 mM PMSF, protease inhibitor cocktail (EDTA-free), 2 mM MgCl_2_, 0.2 mg/mL DNase I) and clarified by centrifugation. Total protein content was measured using the Pierce BCA Protein Assay Kit (Thermo Fisher Scientific, 23225). Equal amounts of protein were separated by SDS–PAGE on self-cast or pre-cast polyacrylamide gels (Bio-Rad, 5678095) and transferred to nitrocellulose membranes (Cytiva, GE10600001) using a wet transfer system. Membranes were blocked in 5% non-fat milk in PBS containing 0.1% Tween-20 (PBS-T) and incubated overnight at 4°C with primary antibodies diluted in the same blocking buffer. Membranes were washed in PBS-T, incubated with HRP-conjugated secondary antibodies for 45 min in blocking solution at room temperature. Chemiluminescence was visualized using Amersham ECL Select Western Blotting Detection Reagent (Cytiva, 12644055) and images acquired with an Alliance LD2 Imaging System (UviTec Cambridge). The following antibodies were used: rat monoclonal anti-Caspase-2 (a gift from Dr. Andreas Strasser, Walter and Eliza Hall Institute of Medical Research, clone 11B4, 1:1000), mouse monoclonal anti-MDM2 (Thermo Fisher Scientific, MA1-113, clone IF2, 1:1000), rabbit monoclonal anti-V5-tag (CST®, 13202, clone D3H8Q, 1:1000), rabbit polyclonal anti-p53 (CST®, 9282, 1:1000), mouse monoclonal anti-MCL1 (SCBT, sc-12756, clone 22, 1:500), rabbit polyclonal anti-CEP83 (Atlas antibodies, HPA038161, 1:1000), rabbit polyclonal anti-PARP1 (CST®, 9542, 1:1000), rabbit polyclonal anti-Caspase-3 (CST®, 9662, 1:1000), rabbit polyclonal anti-Caspase-7 (CST®, 9492, 1:1000), mouse monoclonal anti-CDC27 (BD Biosciences, 610455, clone 35/CDC27, 1:1000), mouse monoclonal anti-p21 (BD Biosciences 564262, clone 2G12, 1:1000), rabbit monoclonal anti-METTL3 (CST®, 96391, clone D2I6O, 1:1000), rabbit polyclonal anti-METTL14 (ProteinTech®, 26158-1-AP, 1:1000), rabbit polyclonal anti-CRADD/RAIDD (ProteinTech®, 10401-1-AP, 1:1000), rabbit monoclonal anti-VIRMA (CST®, 88358, clone D4N8B, 1:1000), rabbit polyclonal anti-WTAP (CST®, 56501, 1:1000), rabbit polyclonal anti-ZC3H13 (Boster Bio, A11022-1, 1:500), rabbit polyclonal anti-CBLL1 (Bethyl Laboratories®, A302-968A, 1:1000), mouse monoclonal anti-RBM15 (ProteinTech®, 66059-1-Ig, clone 4A1A4, 1:1000), mouse monoclonal anti-HSP90 (SCBT, sc-13119, clone F-8, 1:10000), rabbit monoclonal anti-GAPDH (CST®, 2118, clone 14C10, 1:10000), polyclonal goat anti-rat IgG/HRP (Thermo Fisher Scientific, 31470, 1:5000), polyclonal goat anti-rabbit IgG/HRP (Agilent, P0448, 1:5000), polyclonal rabbit anti-mouse IgG/HRP (Agilent, P0161, 1:5000).

### Total RNA extraction and mRNA isolation

Total RNA was isolated using the NucleoSpin RNA Plus Kit (Macherey-Nagel, 740984.50) according to the manufacturer’s instructions.

Polyadenylated mRNA was isolated from total RNA using Dynabeads™ magnetic oligo(dT)25 (Invitrogen, 61005) according to the manufacturer’s guidelines. Dynabeads™ (0.5 mg per sample) were washed once with an equal volume of Binding Buffer (20 mM Tris-HCl - pH 7.5, 1.0 M LiCl, 2 mM EDTA), and resuspended in Binding Buffer to the original beads volume. 100 μL of total RNA (35 μg) in Elution Buffer (10 mM Tris-HCl - pH 7.5) were mixed with 100 μL of Binding Buffer. RNA samples were heated at 65°C for 2 min and immediately placed on ice. The obtained RNA solution was added to washed beads and incubated for 10 min at room temperature under continuous rotation to allow hybridization. Beads were washed once with 200 μL Washing Buffer B (10 mM Tris-HCl - pH 7.5, 0.15 M LiCl, 1 mM EDTA) andpolyadenylated RNA was eluted by resuspending beads in 15 μL Elution Buffer followed by incubation at 80°C for 2 min.

### RT-qPCR

Total RNA was reverse-transcribed using the RevertAid First Strand cDNA Synthesis Kit (Thermo Fisher Scientific, K1622) and random hexamer primers, according to the manufacturer’s instructions. Quantitative PCR was carried out using qPCRBIO Probe Mix No-ROX (Cat# PB20.23), and the 6‐FAM/ZEN/IBFQ dye/quencher-conjugated probes listed in Supplementary Table S1. Amplification was carried out on a Bio-Rad CFX Real-Time PCR System Ct values were obtained using the Bio‐Rad CFX Manager software (version 3.1). Relative gene expression was calculated using the 2^^−ΔΔCt^ method. Mean C_t_ values were derived from technical replicates, normalized to the housekeeping gene to obtain ΔC_t_ values, and subsequently normalized to the untreated wild-type control condition to generate ΔΔC_t_ values.

### ELISA

m6A levels in isolated polyadenylated mRNA were quantified using the m6A RNA Methylation Quantification Kit (Colorimetric) (Abcam, Ab185912) according to the manufacturer’s instructions. Briefly, Binding Solution was first added to each well of the assay plate, followed by 2 μL of Negative Control (NC), 2 μL of diluted Positive Control (PC; 0.5 ng/µL), or 50 ng of sample mRNA. After performing RNA binding, samples were incubated in sequence with Capture Antibody, Detection Antibody, and Enhancer Solution, with washing using Wash Buffer performed between each step. Developer Solution was then added for colorimetric detection, and the reaction was blocked with Stop Solution once the Positive Control developed a medium-blue color. Absorbance (OD) was measured at 450 nm using a Varioskan LUX Microplate Reader (Thermo Fisher Scientific). Relative m6A levels were calculated according to the manufacturer’s formula: m6A (%) = [(OD_sample − OD_NC) / S] ÷ [(OD_PC − OD_NC) / P] × 100, where S represents the input sample RNA amount (ng) and P represents the input Positive Control RNA amount (ng). Values are normalized to the untreated condition.

### Cell sorting and flow cytometry analysis

For cell sorting, cells were harvested by trypsinization, washed with PBS, resuspended in sorting buffer (2 mM EDTA, 1 % BSA in PBS for experiment in Fig. 1E, S1G; 1.5 mM EDTA, 2 % BSA in DMEM/PBS 1:1 for experiment in Fig. 5A) and sorted using a BD FACS Aria III (BD Biosciences) instrument. The obtained cell populations were then subjected to either total protein or genomic DNA isolation for further experiments.

For flow cytometry analysis, cells were harvested by trypsinization, washed with PBS and analyzed with a BD Symphony A1 (BD Biosciences) flow cytometer. Live cells were identified based on forward scatter (FSC) and side scatter (SSC) parameters, and doublets were excluded from the analysis. Analyses of flow cytometry experiments were carried out using FlowJo software (FlowJo, LLC).

### Transcriptomic Library Preparation and analysis

Transcriptomic libraries, including both input RNA-seq and parallel m6A immunoprecipitation fractions, were generated as follows. Total RNA was extracted and poly(A)-enriched prior to fragmentation. Fragmented RNA was barcoded and subjected to immunoprecipitation using an anti-m6A antibody (Synaptic Systems, 202-111), while an aliquot of fragmented RNA was retained as input control. Input and immunoprecipitated fractions were reverse-transcribed, adapter-ligated, PCR-amplified, and quality-controlled according to established procedures^81^. Paired-end libraries were sequenced on an Illumina NovaSeq 6000 platform. Although m6A immunoprecipitation libraries were successfully generated, these experiments did not reveal reproducible enrichment patterns under the conditions tested and were therefore not pursued further. RNA-seq experiments were performed in biological triplicate. Upon FASTQ files demultiplexing (based on perfect matches between reads and sample barcodes - custom R script), reads were aligned to the hg38 genome with STAR using default parameters. Uniquely mapped reads were retained by filtering alignments based on mapping quality (samtools view -hb -q 255). Gene expression levels were quantified from Input samples based on the overlap of both R1 and R2 reads with a unique transcriptional unit (Ensembl annotation 111; R script). Quality control metrics and principal component analysis revealed that one replicate contained shallow sequenced conditions which segregated from the other samples counterparts. This replicate was therefore excluded from downstream differential expression analysis. All reported results are based on the two remaining biological replicates. Exclusion criteria were defined prior to differential expression testing. DESeq2^82^ was used for differential gene expression analysis (default parameters; adjusted *p*-value < 0.1) and log_2_ fold changes shrinkage.

### High-content imaging and kinome-wide screening

For high-content imaging experiments using Cal51^*mScarlet-MDM2*^ cells, caspase/METTL3 inhibitors were administered into 384-well black optically clear flat-bottom plates (Revvity, PhenoPlate 6057302) using the Beckman Coulter Echo 650 (Beckman Coulter, Indianapolis, IN, USA) acoustic liquid handling system. Cellular suspensions treated with either 2 µM ZM447439 or DMSO were seeded directly onto the drug-coated plates using the Integra Viafill multidrop dispenser (INTEGRA Biosciences, Zizers, Swiss) at a density of 5,500 cells/well. Cells were incubated at 37°C for 40 h, then fixed in 4% formaldehyde and nuclei were stained with Hoechst 33342 (1 µg/mL) using the BioTek EL406 Microplate Washer Dispenser (BioTek Instruments Inc., Winooski, VT, USA). Images were acquired using the ImageXpress Micro-Confocal Molecular Devices (3 FOV in widefield mode, 20X Plan Apo Lambda 0.75 NA objective; filter settings: λ_exc_ = 377/54 nm - λ_em_ = 432/36 nm for Hoechst signal and λ_exc_ = 576/23 nm - λ_em_ = 624/40 for mScarlet signal). Similarly, high-content imaging of caspase inhibitors in Cal51 WT cells was performed by administering the caspase inhibitors and 5 µM CellEvent™ Caspase-3/7 Detection Reagent (Thermo Fisher Scientific, C10423) into clear, sterile, tissue culture-treated 384-well plates (Revvity, SpectraPlate 6007650) using the Beckman Coulter Echo 650. Cells were seeded directly onto the drug-coated plates using the Integra Viafill multidrop dispenser at a density of 10,000 cells/well and incubated at 37°C for 16 h. 1 µM staurosporine (STS), 1 µM ABT-737, the combined treatment (STS + ABT) or DMSO were then administered to the cells using Beckman Coulter Echo 650. Cells were incubated at 37°C for additional 4 h and then fixed in 4% formaldehyde and stained with Hoechst 33342 (1 µg/mL) using the BioTek EL406 Microplate Washer Dispenser. Images of the wells were acquired using the ImageXpressMicro-Confocal Molecular Devices (3 FOV in widefield mode, 20X Ph1 Plan Fluor ELWD ADM 0.45NA objective; filter settings: λ_exc_ = 377/54 nm - λ_em_ = 432/36 nm for Hoechst signal and λ_exc_ = 475/28 nm - λ_em_ = 536/40 for CellEvent™ signal). Caspase inhibitors were tested across a 7-point dilution series (0.3125–20 μM), whereas METTL3 inhibitors were tested across a 13-point dilution series (0.0049-20 μM). Data were expressed as percent of positive cells, normalized to the positive control (ZM + DMSO for mScarlet; STS + ABT + DMSO for CellEvent), defined as 100% response. Technical triplicates were averaged prior to analysis, yielding three biological replicate values per concentration. IC_50_values were calculated in GraphPad Prism using nonlinear regression (‘[Inhibitor] vs. response – Variable slope (four parameters)’ model) with comparison to a straight line to distinguish sigmoidal dose-response from inactivity. No additional data transformation was applied. Reported IC_50_values represent means across biological replicates. The KCGS library (Kinase-Chemogenomic Set Library v1.0, provided by the Structural Genomics Consortium (SGC)) screening was performed as described for high-content imaging in Cal51^*mScarlet-MDM2*^ cells (see above) with few exceptions: the used plate types (clear, sterile, tissue culture-treated 384-well plates, Thermo Scientific™, 164688) and the IXM settings (3 FOV in Widefield mode, 20X Ph1 Plan Fluor ELWD ADM 0.45NA objective; filter settings: λ_exc_ = 377/54 nm - λ_em_ = 432/36 nm for Hoechst signal and λ_exc_ = 555/28 nm - λ_em_ = 600/37 for mScarlet signal). KCGS library compounds were tested at 1 µM in duplicate. HTS data were expressed as the duplicate average of Z-score values of Total Cell Count (determined counting Hoechst signals of nuclei) and Positive Nuclei % (percentage of mScarlet-positive nuclei over the total number of nuclei).

### Caspase activities and kinetic analysis

Caspase activities were measured by monitoring the time-dependent hydrolysis of fluorogenic substrates (λ_exc_ = 360 nm; λ_em_ = 460 nm) in 96-well plates using a Fluostar microplate reader at 37 °C. Caspase-2 (0.1 nM, Enzo Life Sciences, ALX-201-057) assays were conducted in buffer containing 20 mM HEPES, 0.1% CHAPS, 5 mM DTT, 2 mM EDTA, 800 mM sodium succinate pH 7.4, and 25 µM Ac-VDVAD-AMC (Enzo Life Sciences, ALX-260-060). Caspase-3 (0.05 nM, R&D Systems, #707-C3) assays were performed in 20 mM HEPES, 0.1% CHAPS, 5 mM DTT, 2 mM EDTA, 1 mg/mL BSA pH 7.4, and 10 µM Ac-DEVD-AMC (Enzo Life Sciences, ALX-260-031). Time-dependent inhibition was analyzed using the continuous progress-curve method^42,83^. Briefly, fluorescence was recorded continuously up to 90 min across a range of inhibitor concentrations. Product formation (in relative fluorescence units, RFU) was plotted as a function of time and fitted to a model describing an initial transient phase, reflecting progressive inactivation, followed by a steady-state phase. The model includes an apparent first-order rate constant (π) that depends on the effective inhibitor concentration [I]’ corrected for substrate competition (using the corresponding K_M_ values) and describes the time course of enzyme inactivation. From global fitting of the progress curves, the kinetic parameters k_inact_ (first-order inactivation rate constant) and K_I_ (apparent inhibitor dissociation constant) were determined. The inhibitory efficiency was expressed as the k_inact_/K_I_ ratio. Linear and nonlinear regressions were performed using the KaleidaGraph software.

### Genome-Wide CRISPR Knockout Screen

For the genome-wide CRISPR knockout screen, lentiviral particles encoding the Brunello sgRNA library were generated as described in the Lentivirus generation section, except that HEK 293T cells were transfected with Metafectene (Biontex, T020-2.0) in place of calcium phosphate-mediated transfection.

Cal51^*mScarlet-MDM2*^ cells were transduced with the Brunello lentiviral library at a multiplicity of infection (MOI) of 0.5 to favor single sgRNA integration. Library representation was maintained at ≥ 250 cells per sgRNA throughout the experiment. Puromycin selection (2 µg/mL) was applied for 72 h starting 24 h post-transduction. Six days after transduction, cells were treated with 2 µM ZM447439 for 40 h to induce whole-genome doubling (WGD). Cells were subsequently sorted based on mScarlet fluorescence, and an unsorted fraction was retained as a reference control. The screen was performed in two independent biological replicates. Genomic DNA was extracted from sorted and control cell populations as previously described^84^. The amount of genomic DNA processed for library amplification was calculated to ensure a representation of at least 250 cells per sgRNA, based on library complexity and genomic DNA content per cell. sgRNA sequences were amplified from genomic DNA by PCR using primers flanking the integrated sgRNA cassette (sequences are available in the Supplementary Table S1). PCR cycling conditions were: 30 s at 98°C; 35 cycles of 10 s at 98°C and 50 s at 72°C; followed by a final extension of 5 min at 72°C. PCR products were purified from agarose gels using the NucleoSpin Gel and PCR Clean-up kit (Macherey-Nagel, 740609). Purified amplicons were quantified, pooled, and subjected to next-generation sequencing on an Illumina NovaSeq 6000 platform using an SP flow cell (Illumina) with 100 bp single-end reads. Sequencing reads were trimmed and assigned to sgRNAs using ReCo^85^. Differential sgRNA representation between sorted and control populations was computed using MAGeCK^52^, and gene-level enrichment scores were derived using the robust ranking aggregation method implemented in MAGeCK.

### Live cell imaging

Cells were seeded in μ-Slide 8 well plates (Ibidi, 80826) at a density of 60,000 cells/well in complete DMEM medium without phenol red (Thermo Fisher Scientific, 21063029). The following day, 1 μM SiR-DNA (Spirochrome, SC007) was added for 1 h. Immediately before imaging, cells were treated with ZM447439 or DMSO.

### Immunofluorescence microscopy

Cells cultured on 18-mm glass coverslips were rinsed with PBS and fixed in 4% formaldehyde (Thermo Fisher Scientific, BP531-500) in PBS for 10 min at room temperature (Fig 1C). Following three 5-min washes in PBS, cells were permeabilized with 0.2% Triton X-100 in PBS for 5 min and washed three additional times. For PIDD1 staining (Fig 7B), cells were permeabilized in PTEM buffer (0.2% Triton X‐100, 20 mM PIPES pH 6.8, 1 mM MgCl_2_, 10 mM EGTA) for 2 min at room temperature and then fixed in 4% formaldehyde in PTEM for an additional 10 minutes. For cells in Fig. S5C, S6A-B, absolute ice‐cold methanol was used to fix and permeabilize cells for at least 20 min at −20°C. Cells were then blocked in 3% BSA (Merck, 112018) in PBS for 20 min. Primary antibodies, diluted in blocking solution, were applied for 1 h at room temperature. After three PBS washes, cells were incubated for 45 min in the dark with fluorophore-conjugated secondary antibodies and 1 μg/mL Hoechst 33342 in blocking solution. Coverslips were washed three times in PBS, briefly dipped in water, and mounted onto glass slides using ProLong Gold Antifade Reagent (Thermo Fisher Scientific, P10144). The following antibodies were used: rabbit polyclonal anti-CEP128 (Atlas antibodies, HPA001116, 1:500), mouse monoclonal anti-PIDD1 (Boster Bio, M10708, clone Anto-1, 1:500), rabbit polyclonal anti-Centrin 1 (Proteintech, 12794-1-AP, 1:500), rabbit polyclonal anti-ANKRD26 (GeneTex, GTX128255, 1:800), mouse monoclonal anti-γ-tubulin (Thermo Fisher Scientific, MA1-19421, clone TU-30, 1:1000), goat polyclonal anti-mouse IgG/AlexaFluor488 (Thermo Fisher Scientific, A-11029, 1:1000), goat polyclonal anti-mouse IgG/AlexaFluor555 (Thermo Fisher Scientific, A-21424, 1:1000), goat polyclonal anti-rabbit IgG/AlexaFluor488 (Thermo Fisher Scientific, A-11034, 1:1000), goat polyclonal anti-rabbit IgG/AlexaFluor555 (Thermo Fisher Scientific, A-21429, 1:1000).

### Ultrastructure Expansion Microscopy

Ultrastructure Expansion Microscopy (U-ExM) was performed according to published protocols^50,86^, with modifications as described by Laporte et al. ^87^. Cells were seeded on 12-mm coverslips and incubated for 3 h at 37°C in PBS containing 2% acrylamide (Thermo Fisher Scientific, BP1402-1) and 1.4% formaldehyde. Gelation was carried out for 1 h at 37°C in PBS containing 19% (w/w) sodium acrylate (Santa Cruz Biotechnology, SC-236893), 10% (w/v) acrylamide, 0.1% (w/v) bis-acrylamide (Thermo Fisher Scientific, BP1404-250), 0.5% (v/v) TEMED (Sigma-Aldrich, T9281), and 0.5% (v/v) ammonium persulfate (Thermo Fisher Scientific, BP179-100). Gels were denatured for 90 min at 95°C in denaturation buffer (200 mM SDS, 200 mM NaCl, 50 mM Tris base pH 9.0) and subsequently transferred to double-distilled water to allow initial expansion.

Expanded gels were equilibrated in PBS for 30 min and incubated overnight at room temperature with primary antibodies diluted in 2% BSA under gentle agitation. After three washes in 0.1% PBS-Tween20, gels were incubated for 2 h at 37°C with fluorophore-conjugated secondary antibodies diluted in 2% BSA. Following three additional washes in 0.1% PBS-Tween20, gels were re-expanded in double-distilled water and mounted on 25-mm poly-D-lysine–coated coverslips for imaging. The following antibodies were used: rabbit polyclonal anti-CEP83 (Atlas antibodies, HPA038161, 1:250), mouse monoclonal anti-acetylated tubulin (Sigma-Aldrich®, T7451, clone 6-11B-1, 1:250), rabbit polyclonal anti-CEP128 (Atlas antibodies, HPA001116, 1:250), goat polyclonal anti-mouse IgG/AlexaFluor555 (Thermo Fisher Scientific, A-21424, 1:500), goat polyclonal anti-rabbit IgG/AlexaFluor488 (Thermo Fisher Scientific, A-11034, 1:500).

### Image acquisition

For live cell imaging experiments, imaging was performed at 37 °C with 5% CO2 on a Nikon Eclipse Ti2E inverted microscope (Nikon Instruments Inc), equipped with Lumencor Spectra X light engine as LED light source, using a plan apochromatic 20x/0.75 dry objective. Images were acquired in widefield mode by an Andor Zyla 4.2 PLUS sCMOS Monochromatic Camera. Single-plane movies were recorded every 5 min for 48 h. For immunofluorescence microscopy and quantification of U-ExM experiments, images were acquired by using a Nikon Eclipse Ti2E inverted microscope (Nikon Instruments Inc) equipped with a Lumencor CELESTA solid-state laser light source, a CrestOptics X-light V3 Spinning Disc module, and an Andor Zyla 4.2 PLUS sCMOS monochromatic camera. The following objectives were used: a plan apochromatic λD 60x/1.42 oil and a plan apochromatic λ 100x/1.45 oil objective. For representative pictures of immunofluorescence experiments, deconvolution was performed with Huygens software (Scientific Volume Imaging, Hilversum, The Netherlands) via Huygens Remote Manager (version 3.10).

For representative pictures of U-ExM experiments, images were acquired with a Nikon Eclipse Ti2E inverted microscope (Nikon Instruments Inc), equipped with a Nikon LU-NV laser light source, a Nikon N-SIM S super resolution structured illumination system, a Nano-Drive MCL piezo electric stage, a Hamamatsu ORCA-Flash 4.0 V3 sCMOS monochromatic camera and a CFI apochromatic SR HP TIRF 100XC/1.49 oil objective. After acquisition, image stacks were reconstructed using the “Reconstruct Stack” function of the NIS-Elements software (version 5.42.06) by applying the same values of “Illumination Modulation Contrast” and “High Resolution Noise Suppression” to all channels.

### Image quantification

All samples stained with the same antibody combinations were acquired using identical imaging settings and processed using consistent intensity scaling. Single-cell nuclear fluorescence of the mScarlet signal was quantified from time-lapse sequences using a custom Fiji macro. Images were uniformly denoised and subjected to background subtraction. The SiR-DNA channel was used both to identify the onset of pseudo-anaphase and to segment individual nuclei. In the resulting binary images, 6-connected components were identified and filtered using the MorphoLibJ plugin to ensure accurate tracking of nuclei throughout the time-lapse. These segmented regions of interest (ROIs) were used to quantify nuclear mScarlet fluorescence over time. Each fluorescence trajectory was aligned to the onset of pseudo-anaphase, defined as time zero for each cell. For immunofluorescence microscopy experiments, PIDD1 fluorescence intensity (Figures 7B and S7A) was quantified by placing circular ROIs (20-pixel diameter) centered on the parent centriole. Signal intensity was measured in Fiji from maximum-intensity projections of z-stacks. For each ROI, background fluorescence -measured in a proximal area lacking centrosomal structures-was subtracted. Fluorescence values were normalized to the mean intensity obtained in control samples.

For characterization of centrosome and appendage integrity (Figure S5D and S6A– C), visual scoring was performed on conventional immunofluorescence images. Centriole numbers were counted based on γ-tubulin signal, distal appendages were scored as present or absent based on detection of CEP83 signal at mother centrioles and subdistal appendage integrity was assessed based on the presence or absence of CEP128 staining at centrosomes. For U-ExM experiments, gel expansion factors were calculated by measuring gel diameters after expansion using a caliper and dividing these values by the initial gel diameter prior denaturation (12 mm). The resulting expansion factors were used to rescale scale bars and all reported measurements accordingly. For quantification of centrosome clustering, the three-dimensional distance between the two closest CEP83-positive signals was manually measured in Fiji by selecting the point of maximal intensity within each signal.

### Statistical analysis

Over-representation analysis (ORA) and gene set enrichment analysis (GSEA) were performed in R (version 4.5.2) using the clusterProfiler package. ORA for Gene Ontology was carried out on the top 200 CRISPR screen hits, using the Brunello library as the background universe and applying Benjamini–Hochberg correction; terms with p adj < 0.05 and q < 0.2 were retained. The top five enriched GO Cellular component (CC) terms were visualized as a dot plot, and their associated gene sets were used to calculate pairwise Jaccard similarity coefficients, which were displayed as a clustered heatmap. GSEA was performed on all screen genes ranked by MAGeCK score against GO CC terms restricted to genes present in the Brunello background. The analysis was run with a minimum gene set size of 10, maximum of 500, and a p-value cutoff of 0.05 using positive enrichment scores. Enrichment plots for the GO terms “centriole” and “subdistal appendage” are shown in the figures. Data are presented as dot plots, Superplots^88^ ,bar charts or box plots, showing the mean ± standard deviation unless otherwise specified in the corresponding figure legend. In dot plots, different biological replicates are color-coded. Normality of datasets was assessed using the Shapiro–Wilk test. Statistical differences were evaluated using unpaired two‐tailed Student’s *t*‐test or Mann–Whitney test for comparison between two groups, and one‐way ANOVA or Kruskal–Wallis test for comparison among multiple groups, with Tukey’s or Dunn’s multiple comparisons test applied, respectively. Statistical analyses and graphs generation were performed using GraphPad Prism 10 (GraphPad, San Diego, CA, USA).

## Acknowledgments

We thank Drs. Alessio Zippo for critical reading of the manuscript, Alessandro Provenzani and Adrew Duncan for useful discussions, Laura Alunno and Vincenza Vigorito for laboratory assistance, and the CIBIO Core Facilities and their staff for expert technical assistance. This work was supported by AIRC (MFAG, project ID 23560 to L.L.F; Bridge Grant, project ID 30372 to L.L.F.; IG, project ID 31900 to L.L.F.; IG, project ID 22075 to A.Q.; IG, project ID 25849 to A.I.), Telethon (GJC21181 to L.L.F.), the Giovanni Armenise‐Harvard Foundation (CDA 2017 to L.L.F.), the University of Trento (to L.L.F., D.M., and N.C.), the European Union - NextGenerationEU (PNRR M4C2 INV 1.1, PRIN 2022 PNRR Prot. n. P2022A9J9L and PRIN 2022 Prot. n. 2022L8RAKN to L.L.F.) and the initiative “Dipartimenti di Eccellenza 2023-2027 (Legge 232/2016)” funded by the MUR. D.M. was supported by an AIRC fellowship for Italy (ID 32875-2025). Department CIBIO Core Facilities (IRBIO) are supported by the European Regional Development Fund (ERDF) 2014-2020 and 2021-2027. P.G., V.H. and M.H.L. were supported by the Swiss National Science Foundation (SNSF) 310030_205087.

## Author Contributions

Conceptualization, L.L.F., and M.B.; Methodology, S.T., M.Pa., M.F., L.C., M.La, M.Li, S.B., A.I., V.H., P.G., M.Pe., and M.K.; Investigation, D.M., A.M., G.M.M., N.C., G.P., L.C., F.S., and M.B.; Formal Analysis, D.M., M.Pa., M.F., M.W., T.T., and M.B.; Resources, L.L.F., A.Q., E.D.J., V.H., and P.G.; Data Curation, D.M., M.F., and M.B.; Writing – Original Draft, L.L.F., M.B., and D.M.; Writing – Review & Editing, all authors; Supervision, L.L.F., M.B., and D.M.; Funding Acquisition, L.L.F.

## Declaration of generative AI and AI-assisted technologies

During the preparation of this manuscript, the authors used ChatGPT to enhance the writing style. All content generated with the assistance of the tool was thoroughly reviewed and edited by the authors, who take full responsibility for the final version of the publication.

## References

1. Fujiwara, T., Bandi, M., Nitta, M., Ivanova, E.V., Bronson, R.T., and Pellman, D. (2005). Cytokinesis failure generating tetraploids promotes tumorigenesis in p53-null cells. Nature 437, 1043–1047. 10.1038/nature04217.

2. Ganem, N.J., Godinho, S.A., and Pellman, D. (2009). A mechanism linking extra centrosomes to chromosomal instability. Nature 460, 278–282. 10.1038/nature08136.

3. Gemble, S., Wardenaar, R., Keuper, K., Srivastava, N., Nano, M., Mace, A.S., Tijhuis, A.E., Bernhard, S.V., Spierings, D.C.J., Simon, A., et al. (2022). Genetic instability from a single S phase after whole-genome duplication. Nature 604, 146–151. 10.1038/s41586-022-04578-4.

4. Dewhurst, S.M., McGranahan, N., Burrell, R.A., Rowan, A.J., Gronroos, E., Endesfelder, D., Joshi, T., Mouradov, D., Gibbs, P., Ward, R.L., et al. (2014). Tolerance of whole-genome doubling propagates chromosomal instability and accelerates cancer genome evolution. Cancer Discov 4, 175–185. 10.1158/2159-8290.CD-13-0285.

5. Vittoria, M.A., Quinton, R.J., and Ganem, N.J. (2023). Whole-genome doubling in tissues and tumors. Trends Genet 39, 954–967. 10.1016/j.tig.2023.08.004.

6. Zack, T.I., Schumacher, S.E., Carter, S.L., Cherniack, A.D., Saksena, G., Tabak, B., Lawrence, M.S., Zhang, C.Z., Wala, J., Mermel, C.H., et al. (2013). Pan-cancer patterns of somatic copy number alteration. Nat Genet 45, 1134–1140. 10.1038/ng.2760.

7. Bielski, C.M., Zehir, A., Penson, A.V., Donoghue, M.T.A., Chatila, W., Armenia, J., Chang, M.T., Schram, A.M., Jonsson, P., Bandlamudi, C., et al. (2018). Genome doubling shapes the evolution and prognosis of advanced cancers. Nat Genet 50, 1189–1195. 10.1038/s41588-018-0165-1.

8. Minussi, D.C., Nicholson, M.D., Ye, H., Davis, A., Wang, K., Baker, T., Tarabichi, M., Sei, E., Du, H., Rabbani, M., et al. (2021). Breast tumours maintain a reservoir of subclonal diversity during expansion. Nature 592, 302–308. 10.1038/s41586-021-03357-x.

9. Quinton, R.J., DiDomizio, A., Vittoria, M.A., Kotynkova, K., Ticas, C.J., Patel, S., Koga, Y., Vakhshoorzadeh, J., Hermance, N., Kuroda, T.S., et al. (2021). Whole-genome doubling confers unique genetic vulnerabilities on tumour cells. Nature 590, 492–497. 10.1038/s41586-020-03133-3.

10. Aylon, Y., and Oren, M. (2011). p53: guardian of ploidy. Mol Oncol 5, 315–323. 10.1016/j.molonc.2011.07.007.

11. Ganem, N.J., and Pellman, D. (2007). Limiting the proliferation of polyploid cells. Cell 131, 437–440. 10.1016/j.cell.2007.10.024.

12. Margolis, R.L., Lohez, O.D., and Andreassen, P.R. (2003). G1 tetraploidy checkpoint and the suppression of tumorigenesis. J Cell Biochem 88, 673–683. 10.1002/jcb.10411.

13. Wright, W.E., and Hayflick, L. (1972). Formation of anucleate and multinucleate cells in normal and SV 40 transformed WI-38 by cytochalasin B. Exp Cell Res 74, 187–194. 10.1016/0014-4827(72)90496-x.

14. Fava, L.L., Schuler, F., Sladky, V., Haschka, M.D., Soratroi, C., Eiterer, L., Demetz, E., Weiss, G., Geley, S., Nigg, E.A., and Villunger, A. (2017). The PIDDosome activates p53 in response to supernumerary centrosomes. Genes Dev 31, 34–45. 10.1101/gad.289728.116.

15. Ganem, N.J., Cornils, H., Chiu, S.Y., O’Rourke, K.P., Arnaud, J., Yimlamai, D., Thery, M., Camargo, F.D., and Pellman, D. (2014). Cytokinesis failure triggers hippo tumor suppressor pathway activation. Cell 158, 833–848. 10.1016/j.cell.2014.06.029.

16. Burigotto, M., Mattivi, A., Migliorati, D., Magnani, G., Valentini, C., Roccuzzo, M., Offterdinger, M., Pizzato, M., Schmidt, A., Villunger, A., et al. (2021). Centriolar distal appendages activate the centrosome-PIDDosome-p53 signalling axis via ANKRD26. EMBO J 40, e104844. 10.15252/embj.2020104844.

17. Eichin, F., Sladky, V.C., Reiner, M.A., Leone, M., Abila, E., Rendeiro, A.F., Böttcher, R., Dahlhoff, M., Kolbe, T., and Villunger, A. (2025). 10.1101/2025.06.23.660994.

18. Evans, L.T., Anglen, T., Scott, P., Lukasik, K., Loncarek, J., and Holland, A.J. (2021). ANKRD26 recruits PIDD1 to centriolar distal appendages to activate the PIDDosome following centrosome amplification. EMBO J 40, e105106. 10.15252/embj.2020105106.

19. Leone, M., Kinz, N., Eichin, F., Obwegs, D., Sladky, V.C., Braun, V.Z., Hirschberger, R., Rizzotto, D., Englmaier, L., Manzl, C., et al. (2026). The PIDDosome controls cardiomyocyte polyploidization during postnatal heart development. Cell Death Differ. 10.1038/s41418-025-01645-x.

20. Rizzotto, D., Vigorito, V., Rieder, P., Gallob, F., Moretta, G.M., Soratroi, C., Riley, J.S., Bellutti, F., Veli, S.L., Mattivi, A., et al. (2024). Caspase-2 kills cells with extra centrosomes. Sci Adv 10, eado6607. 10.1126/sciadv.ado6607.

21. Sladky, V.C., Akbari, H., Tapias-Gomez, D., Evans, L.T., Drown, C.G., Strong, M.A., LoMastro, G.M., Larman, T., and Holland, A.J. (2022). Centriole signaling restricts hepatocyte ploidy to maintain liver integrity. Genes Dev 36, 843–856. 10.1101/gad.349727.122.

22. Sladky, V.C., Knapp, K., Soratroi, C., Heppke, J., Eichin, F., Rocamora-Reverte, L., Szabo, T.G., Bongiovanni, L., Westendorp, B., Moreno, E., et al. (2020). E2F-Family Members Engage the PIDDosome to Limit Hepatocyte Ploidy in Liver Development and Regeneration. Dev Cell 52, 335–349 e337. 10.1016/j.devcel.2019.12.016.

23. Kopeina, G.S., and Zhivotovsky, B. (2021). Caspase-2 as a master regulator of genomic stability. Trends Cell Biol 31, 712–720. 10.1016/j.tcb.2021.03.002.

24. Tinel, A., and Tschopp, J. (2004). The PIDDosome, a protein complex implicated in activation of caspase-2 in response to genotoxic stress. Science 304, 843–846. 10.1126/science.1095432.

25. Oliver, T.G., Meylan, E., Chang, G.P., Xue, W., Burke, J.R., Humpton, T.J., Hubbard, D., Bhutkar, A., and Jacks, T. (2011). Caspase-2-mediated cleavage of Mdm2 creates a p53-induced positive feedback loop. Mol Cell 43, 57–71. 10.1016/j.molcel.2011.06.012.

26. Talanian, R.V., Quinlan, C., Trautz, S., Hackett, M.C., Mankovich, J.A., Banach, D., Ghayur, T., Brady, K.D., and Wong, W.W. (1997). Substrate specificities of caspase family proteases. J Biol Chem 272, 9677–9682. 10.1074/jbc.272.15.9677.

27. Wejda, M., Impens, F., Takahashi, N., Van Damme, P., Gevaert, K., and Vandenabeele, P. (2012). Degradomics reveals that cleavage specificity profiles of caspase-2 and effector caspases are alike. J Biol Chem 287, 33983–33995. 10.1074/jbc.M112.384552.

28. Ando, K., Parsons, M.J., Shah, R.B., Charendoff, C.I., Paris, S.L., Liu, P.H., Fassio, S.R., Rohrman, B.A., Thompson, R., Oberst, A., et al. (2017). NPM1 directs PIDDosome-dependent caspase-2 activation in the nucleolus. J Cell Biol 216, 1795–1810. 10.1083/jcb.201608095.

29. Bouchier-Hayes, L., Oberst, A., McStay, G.P., Connell, S., Tait, S.W., Dillon, C.P., Flanagan, J.M., Beere, H.M., and Green, D.R. (2009). Characterization of cytoplasmic caspase-2 activation by induced proximity. Mol Cell 35, 830–840. 10.1016/j.molcel.2009.07.023.

30. Shah, R.B., Li, Y., Yu, H., Kini, E., and Sidi, S. (2024). Stepwise phosphorylation and SUMOylation of PIDD1 drive PIDDosome assembly in response to DNA repair failure. Nat Commun 15, 9195. 10.1038/s41467-024-53412-0.

31. Barbieri, I., and Kouzarides, T. (2020). Role of RNA modifications in cancer. Nat Rev Cancer 20, 303–322. 10.1038/s41568-020-0253-2.

32. Deng, X., Wu, D., Zhao, Y., Qing, Y., Wu, H., and Chen, J. (2025). Epitranscriptomic control of cancer hallmarks: Functions, mechanisms, and therapeutics of RNA modifications. Cancer Cell. 10.1016/j.ccell.2025.12.001.

33. Luo, H., Kharas, M.G., and Jaffrey, S.R. (2026). N(6)-Methyladenosine: an RNA modification as a central regulator of cancer. Nat Rev Cancer 26, 118–136. 10.1038/s41568-025-00889-6.

34. Elvira-Blazquez, D., Fernandez-Justel, J.M., Arcas, A., Statello, L., Goni, E., Gonzalez, J., Ricci, B., Zaccara, S., Raimondi, I., and Huarte, M. (2024). YTHDC1 m(6)A-dependent and m(6)A-independent functions converge to preserve the DNA damage response. EMBO J 43, 3494–3522. 10.1038/s44318-024-00153-x.

35. Raj, N., Wang, M., Seoane, J.A., Zhao, R.L., Kaiser, A.M., Moonie, N.A., Demeter, J., Boutelle, A.M., Kerr, C.H., Mulligan, A.S., et al. (2022). The Mettl3 epitranscriptomic writer amplifies p53 stress responses. Mol Cell 82, 2370–2384 e2310. 10.1016/j.molcel.2022.04.010.

36. Zhang, J., Wei, J., Sun, R., Sheng, H., Yin, K., Pan, Y., Jimenez, R., Chen, S., Cui, X.L., Zou, Z., et al. (2023). A lncRNA from the FTO locus acts as a suppressor of the m(6)A writer complex and p53 tumor suppression signaling. Mol Cell 83, 2692–2708 e2697. 10.1016/j.molcel.2023.06.024.

37. Ditchfield, C., Johnson, V.L., Tighe, A., Ellston, R., Haworth, C., Johnson, T., Mortlock, A., Keen, N., and Taylor, S.S. (2003). Aurora B couples chromosome alignment with anaphase by targeting BubR1, Mad2, and Cenp-E to kinetochores. J Cell Biol 161, 267–280. 10.1083/jcb.200208091jcb.200208091 [pii].

38. Bindels, D.S., Haarbosch, L., van Weeren, L., Postma, M., Wiese, K.E., Mastop, M., Aumonier, S., Gotthard, G., Royant, A., Hink, M.A., and Gadella, T.W., Jr. (2017). mScarlet: a bright monomeric red fluorescent protein for cellular imaging. Nat Methods 14, 53–56. 10.1038/nmeth.4074.

39. Linton, S.D., Aja, T., Armstrong, R.A., Bai, X., Chen, L.S., Chen, N., Ching, B., Contreras, P., Diaz, J.L., Fisher, C.D., et al. (2005). First-in-class pan caspase inhibitor developed for the treatment of liver disease. J Med Chem 48, 6779–6782. 10.1021/jm050307e.

40. Caserta, T.M., Smith, A.N., Gultice, A.D., Reedy, M.A., and Brown, T.L. (2003). Q-VD-OPh, a broad spectrum caspase inhibitor with potent antiapoptotic properties. Apoptosis 8, 345–352. 10.1023/a:1024116916932.

41. Chauvier, D., Ankri, S., Charriaut-Marlangue, C., Casimir, R., and Jacotot, E. (2007). Broad-spectrum caspase inhibitors: from myth to reality? Cell Death Differ 14, 387–391. 10.1038/sj.cdd.4402044.

42. Bosc, E., Anastasie, J., Soualmia, F., Coric, P., Kim, J.Y., Wang, L.Q., Lacin, G., Zhao, K., Patel, R., Duplus, E., et al. (2022). Genuine selective caspase-2 inhibition with new irreversible small peptidomimetics. Cell Death Dis 13, 959. 10.1038/s41419-022-05396-2.

43. Wannamaker, W., Davies, R., Namchuk, M., Pollard, J., Ford, P., Ku, G., Decker, C., Charifson, P., Weber, P., Germann, U.A., et al. (2007). (S)-1-((S)-2-[1-(4-amino-3-chloro-phenyl)-methanoyl]-amino-3,3-dimethyl-butanoyl)-pyrrolidine-2-carboxylic acid ((2R,3S)-2-ethoxy-5-oxo-tetrahydro-furan-3-yl)-amide (VX-765), an orally available selective interleukin (IL)-converting enzyme/caspase-1 inhibitor, exhibits potent anti-inflammatory activities by inhibiting the release of IL-1beta and IL-18. J Pharmacol Exp Ther 321, 509–516. 10.1124/jpet.106.111344.

44. Wells, C.I., Al-Ali, H., Andrews, D.M., Asquith, C.R.M., Axtman, A.D., Dikic, I., Ebner, D., Ettmayer, P., Fischer, C., Frederiksen, M., et al. (2021). The Kinase Chemogenomic Set (KCGS): An Open Science Resource for Kinase Vulnerability Identification. Int J Mol Sci 22. 10.3390/ijms22020566.

45. Steegmaier, M., Hoffmann, M., Baum, A., Lenart, P., Petronczki, M., Krssak, M., Gurtler, U., Garin-Chesa, P., Lieb, S., Quant, J., et al. (2007). BI 2536, a potent and selective inhibitor of polo-like kinase 1, inhibits tumor growth in vivo. Curr Biol 17, 316–322. 10.1016/j.cub.2006.12.037.

46. Burkard, M.E., Randall, C.L., Larochelle, S., Zhang, C., Shokat, K.M., Fisher, R.P., and Jallepalli, P.V. (2007). Chemical genetics reveals the requirement for Polo-like kinase 1 activity in positioning RhoA and triggering cytokinesis in human cells. Proc Natl Acad Sci U S A 104, 4383–4388. 10.1073/pnas.0701140104.

47. Kong, D., Farmer, V., Shukla, A., James, J., Gruskin, R., Kiriyama, S., and Loncarek, J. (2014). Centriole maturation requires regulated Plk1 activity during two consecutive cell cycles. J Cell Biol 206, 855–865. 10.1083/jcb.201407087.

48. Tsou, M.F., Wang, W.J., George, K.A., Uryu, K., Stearns, T., and Jallepalli, P.V. (2009). Polo kinase and separase regulate the mitotic licensing of centriole duplication in human cells. Dev Cell 17, 344–354. 10.1016/j.devcel.2009.07.015.

49. Wang, W.J., Soni, R.K., Uryu, K., and Tsou, M.F. (2011). The conversion of centrioles to centrosomes: essential coupling of duplication with segregation. J Cell Biol 193, 727–739. 10.1083/jcb.201101109.

50. Gambarotto, D., Zwettler, F.U., Le Guennec, M., Schmidt-Cernohorska, M., Fortun, D., Borgers, S., Heine, J., Schloetel, J.G., Reuss, M., Unser, M., et al. (2019). Imaging cellular ultrastructures using expansion microscopy (U-ExM). Nat Methods 16, 71–74. 10.1038/s41592-018-0238-1.

51. Doench, J.G., Fusi, N., Sullender, M., Hegde, M., Vaimberg, E.W., Donovan, K.F., Smith, I., Tothova, Z., Wilen, C., Orchard, R., et al. (2016). Optimized sgRNA design to maximize activity and minimize off-target effects of CRISPR-Cas9. Nat Biotechnol 34, 184–191. 10.1038/nbt.3437.

52. Li, W., Xu, H., Xiao, T., Cong, L., Love, M.I., Zhang, F., Irizarry, R.A., Liu, J.S., Brown, M., and Liu, X.S. (2014). MAGeCK enables robust identification of essential genes from genome-scale CRISPR/Cas9 knockout screens. Genome Biol 15, 554. 10.1186/s13059-014-0554-4.

53. Baugh, E.H., Ke, H., Levine, A.J., Bonneau, R.A., and Chan, C.S. (2018). Why are there hotspot mutations in the TP53 gene in human cancers? Cell Death Differ 25, 154–160. 10.1038/cdd.2017.180.

54. Lin, Y., Ma, W., and Benchimol, S. (2000). Pidd, a new death-domain-containing protein, is induced by p53 and promotes apoptosis. Nat Genet 26, 122–127. 10.1038/79102.

55. Barak, Y., Juven, T., Haffner, R., and Oren, M. (1993). mdm2 expression is induced by wild type p53 activity. EMBO J 12, 461–468. 10.1002/j.1460-2075.1993.tb05678.x.

56. Wang, P., Doxtader, K.A., and Nam, Y. (2016). Structural Basis for Cooperative Function of Mettl3 and Mettl14 Methyltransferases. Mol Cell 63, 306–317. 10.1016/j.molcel.2016.05.041.

57. Dolbois, A., Bedi, R.K., Bochenkova, E., Muller, A., Moroz-Omori, E.V., Huang, D., and Caflisch, A. (2021). 1,4,9-Triazaspiro[5.5]undecan-2-one Derivatives as Potent and Selective METTL3 Inhibitors. J Med Chem 64, 12738–12760. 10.1021/acs.jmedchem.1c00773.

58. Guirguis, A.A., Ofir-Rosenfeld, Y., Knezevic, K., Blackaby, W., Hardick, D., Chan, Y.C., Motazedian, A., Gillespie, A., Vassiliadis, D., Lam, E.Y.N., et al. (2023). Inhibition of METTL3 Results in a Cell-Intrinsic Interferon Response That Enhances Antitumor Immunity. Cancer Discov 13, 2228–2247. 10.1158/2159-8290.CD-23-0007.

59. Moser, J.C., Papadopoulos, K.P., Rodon Ahnert, J., Ofir-Rosenfeld, Y., and Holz, J.-B. (2024). Phase 1 dose escalation and cohort expansion study evaluating safety, PK, PD and clinical activity of STC-15, a METTL-3 inhibitor, in patients with advanced malignancies. Journal of Clinical Oncology 42, 2586–2586. 10.1200/JCO.2024.42.16_suppl.2586.

60. Yankova, E., Blackaby, W., Albertella, M., Rak, J., De Braekeleer, E., Tsagkogeorga, G., Pilka, E.S., Aspris, D., Leggate, D., Hendrick, A.G., et al. (2021). Small-molecule inhibition of METTL3 as a strategy against myeloid leukaemia. Nature 593, 597–601. 10.1038/s41586-021-03536-w.

61. Karasu, O.R., Neuner, A., Atorino, E.S., Pereira, G., and Schiebel, E. (2022). The central scaffold protein CEP350 coordinates centriole length, stability, and maturation. J Cell Biol 221. 10.1083/jcb.202203081.

62. Mahjoub, M.R., Xie, Z., and Stearns, T. (2010). Cep120 is asymmetrically localized to the daughter centriole and is essential for centriole assembly. J Cell Biol 191, 331–346. 10.1083/jcb.201003009.

63. Mazo, G., Soplop, N., Wang, W.J., Uryu, K., and Tsou, M.F. (2016). Spatial Control of Primary Ciliogenesis by Subdistal Appendages Alters Sensation-Associated Properties of Cilia. Dev Cell 39, 424–437. 10.1016/j.devcel.2016.10.006.

64. Pudlowski, R., Xu, L., Milenkovic, L., Kumar, C., Hemsworth, K., Aqrabawi, Z., Stearns, T., and Wang, J.T. (2025). A delta-tubulin/epsilon-tubulin/Ted protein complex is required for centriole architecture. Elife 13. 10.7554/eLife.98704.

65. Tanos, B.E., Yang, H.J., Soni, R., Wang, W.J., Macaluso, F.P., Asara, J.M., and Tsou, M.F. (2013). Centriole distal appendages promote membrane docking, leading to cilia initiation. Genes Dev 27, 163–168. 10.1101/gad.207043.112.

66. Tsai, J.J., Hsu, W.B., Liu, J.H., Chang, C.W., and Tang, T.K. (2019). CEP120 interacts with C2CD3 and Talpid3 and is required for centriole appendage assembly and ciliogenesis. Sci Rep 9, 6037 10.1038/s41598-019-42577-0.

67. Chong, W.M., Wang, W.J., Lo, C.H., Chiu, T.Y., Chang, T.J., Liu, Y.P., Tanos, B., Mazo, G., Tsou, M.B., Jane, W.N., et al. (2020). Super-resolution microscopy reveals coupling between mammalian centriole subdistal appendages and distal appendages. Elife 9. 10.7554/eLife.53580.

68. Tinel, A., Janssens, S., Lippens, S., Cuenin, S., Logette, E., Jaccard, B., Quadroni, M., and Tschopp, J. (2007). Autoproteolysis of PIDD marks the bifurcation between pro-death caspase-2 and pro-survival NF-kappaB pathway. EMBO J 26, 197–208. 10.1038/sj.emboj.7601473.

69. Huang, N., Xia, Y., Zhang, D., Wang, S., Bao, Y., He, R., Teng, J., and Chen, J. (2017). Hierarchical assembly of centriole subdistal appendages via centrosome binding proteins CCDC120 and CCDC68. Nat Commun 8, 15057. 10.1038/ncomms15057.

70. Haupt, Y., Maya, R., Kazaz, A., and Oren, M. (1997). Mdm2 promotes the rapid degradation of p53. Nature 387, 296–299. 10.1038/387296a0.

71. Honda, R., Tanaka, H., and Yasuda, H. (1997). Oncoprotein MDM2 is a ubiquitin ligase E3 for tumor suppressor p53. FEBS Lett 420, 25–27. 10.1016/s0014-5793(97)01480-4.

72. Kubbutat, M.H., Jones, S.N., and Vousden, K.H. (1997). Regulation of p53 stability by Mdm2. Nature 387, 299–303. 10.1038/387299a0.

73. Lahav, G., Rosenfeld, N., Sigal, A., Geva-Zatorsky, N., Levine, A.J., Elowitz, M.B., and Alon, U. (2004). Dynamics of the p53-Mdm2 feedback loop in individual cells. Nat Genet 36, 147–150. 10.1038/ng1293.

74. Purvis, J.E., Karhohs, K.W., Mock, C., Batchelor, E., Loewer, A., and Lahav, G. (2012). p53 dynamics control cell fate. Science 336, 1440–1444. 10.1126/science.1218351.

75. Tsabar, M., Mock, C.S., Venkatachalam, V., Reyes, J., Karhohs, K.W., Oliver, T.G., Regev, A., Jambhekar, A., and Lahav, G. (2020). A Switch in p53 Dynamics Marks Cells That Escape from DSB-Induced Cell Cycle Arrest. Cell Rep 32, 107995. 10.1016/j.celrep.2020.107995.

76. Barajas, J.M., Lin, C.H., Sun, H.L., Alencastro, F., Zhu, A.C., Aljuhani, M., Navari, L., Yilmaz, S.A., Yu, L., Corps, K., et al. (2022). METTL3 Regulates Liver Homeostasis, Hepatocyte Ploidy, and Circadian Rhythm-Controlled Gene Expression in Mice. Am J Pathol 192, 56–71. 10.1016/j.ajpath.2021.09.005.

77. Pizzato, M., Erlwein, O., Bonsall, D., Kaye, S., Muir, D., and McClure, M.O. (2009). A one-step SYBR Green I-based product-enhanced reverse transcriptase assay for the quantitation of retroviruses in cell culture supernatants. J Virol Methods 156, 1–7. 10.1016/j.jviromet.2008.10.012.

78. Ghetti, S., Burigotto, M., Mattivi, A., Magnani, G., Casini, A., Bianchi, A., Cereseto, A., and Fava, L.L. (2021). CRISPR/Cas9 ribonucleoprotein-mediated knockin generation in hTERT-RPE1 cells. STAR Protoc 2, 100407. 10.1016/j.xpro.2021.100407.

79. Conant, D., Hsiau, T., Rossi, N., Oki, J., Maures, T., Waite, K., Yang, J., Joshi, S., Kelso, R., Holden, K., et al. (2022). Inference of CRISPR Edits from Sanger Trace Data. CRISPR J 5, 123–130. 10.1089/crispr.2021.0113.

80. Edelheit, O., Hanukoglu, A., and Hanukoglu, I. (2009). Simple and efficient site-directed mutagenesis using two single-primer reactions in parallel to generate mutants for protein structure-function studies. BMC Biotechnol 9, 61. 10.1186/1472-6750-9-61.

81. Dierks, D., Garcia-Campos, M.A., Uzonyi, A., Safra, M., Edelheit, S., Rossi, A., Sideri, T., Varier, R.A., Brandis, A., Stelzer, Y., et al. (2021). Multiplexed profiling facilitates robust m6A quantification at site, gene and sample resolution. Nat Methods 18, 1060–1067. 10.1038/s41592-021-01242-z.

82. Love, M.I., Huber, W., and Anders, S. (2014). Moderated estimation of fold change and dispersion for RNA-seq data with DESeq2. Genome Biol 15, 550. 10.1186/s13059-014-0550-8.

83. Doucet, C., Pochet, L., Thierry, N., Pirotte, B., Delarge, J., and Reboud-Ravaux, M. (1999). 6-Substituted 2-oxo-2H-1-benzopyran-3-carboxylic acid as a core structure for specific inhibitors of human leukocyte elastase. J Med Chem 42, 4161–4171. 10.1021/jm990070k.

84. Diehl, V., Wegner, M., Grumati, P., Husnjak, K., Schaubeck, S., Gubas, A., Shah, V.J., Polat, I.H., Langschied, F., Prieto-Garcia, C., et al. (2021). Minimized combinatorial CRISPR screens identify genetic interactions in autophagy. Nucleic Acids Res 49, 5684–5704. 10.1093/nar/gkab309.

85. Wegner, M., and Kaulich, M. (2023). ReCo: automated NGS read-counting of single and combinatorial CRISPR gRNAs. Bioinformatics 39. 10.1093/bioinformatics/btad448.

86. Gambarotto, D., Hamel, V., and Guichard, P. (2021). Ultrastructure expansion microscopy (U-ExM). Methods Cell Biol 161, 57–81. 10.1016/bs.mcb.2020.05.006.

87. Laporte, M.H., Bouhlel, I.B., Bertiaux, E., Morrison, C.G., Giroud, A., Borgers, S., Azimzadeh, J., Bornens, M., Guichard, P., Paoletti, A., and Hamel, V. (2022). Human SFI1 and Centrin form a complex critical for centriole architecture and ciliogenesis. EMBO J 41, e112107. 10.15252/embj.2022112107.

88. Lord, S.J., Velle, K.B., Mullins, R.D., and Fritz-Laylin, L.K. (2020). SuperPlots: Communicating reproducibility and variability in cell biology. J Cell Biol 219. 10.1083/jcb.202001064.

