## Supplementary Information for "Centrosome architecture and m6A-dependent gating of p53 surveillance after whole-genome doubling"

Supplementary Information
Related to

"Centrosome architecture and m6A-dependent gating of p53 surveillance after whole-genome doubling"

**Figures S1-S7**
**Supplementary Movie 1, legend**
**Supplementary Table S1**
**Supplementary References**

Migliorati, Mattivi, Moretta et al., Figure S1

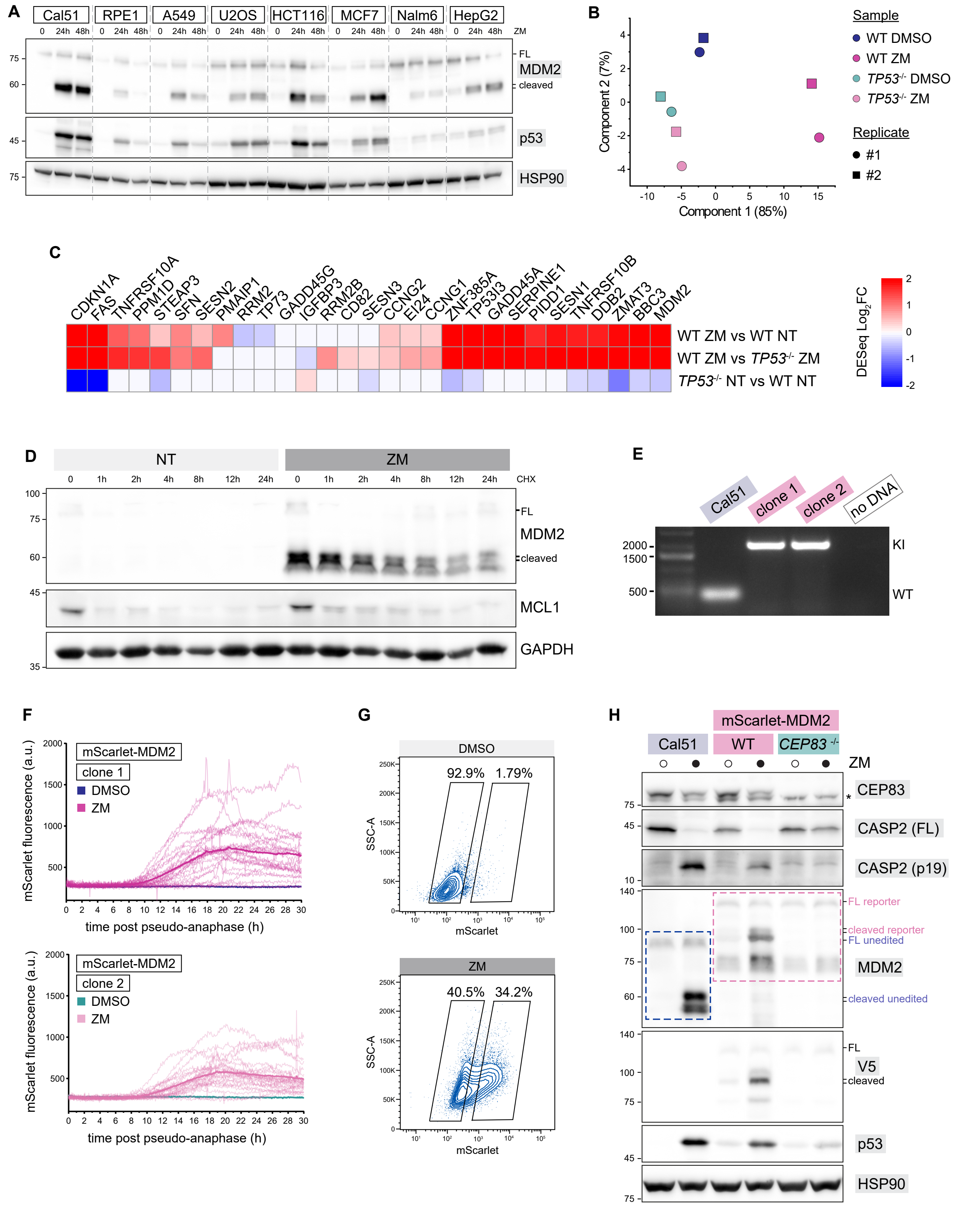


**Figure S1. Selection and validation of Cal51 as a model for Caspase-2 activation and reporter characterization, related to Figure 1**

**(A)** Immunoblot analysis of MDM2 cleavage in a panel of human cell lines untreated or treated with ZM for the indicated times.

**(B)** Principal component analysis (PCA) of bulk RNA-seq profiles from wild-type (WT) and *TP53*$^{-/-}$ Cal51 cells treated with DMSO or ZM. N = 2 independent replicates.

**(C)** Heatmap of representative p53 target genes upregulated in ZM-treated WT Cal51 cells but not in *TP53*$^{-/-}$ cells. Values represent $\log_2$-transformed, Z-score–normalized expression levels across conditions.
**(D)** Cycloheximide (CHX) chase assay in ZM-treated Cal51 cells, followed by immunoblot analysis at the indicated time points.
**(E)** Genotyping PCR of the indicated Cal51$^{mScarlet\text{-}MDM2}$ clones. WT = wild type; KI = knock-in.
**(F)** Individual mScarlet fluorescence traces of Cal51$^{mScarlet\text{-}MDM2}$ cells imaged live after ZM or vehicle-only treatment, corresponding to the quantification shown in Fig. 1D. Traces were aligned by setting time zero to the onset of pseudo-anaphase for each cell. The darker trace represents the population mean (same data as in Fig. 1D), whereas lighter traces show fluorescence profiles from individual cells (n = 25 cells per condition).
**(G)** Flow cytometry analysis of Cal51$^{mScarlet\text{-}MDM2}$ cells treated with ZM, showing the gating strategy used to isolate mScarlet-positive and mScarlet-negative populations for immunoblot analysis.
**(H)** Immunoblot analysis of unedited Cal51, wild-type (WT), and *CEP83*$^{-/-}$ Cal51$^{mScarlet\text{-}MDM2}$ cells treated with ZM for 40 h, corresponding to the flow cytometry profiles shown in Fig. 1F

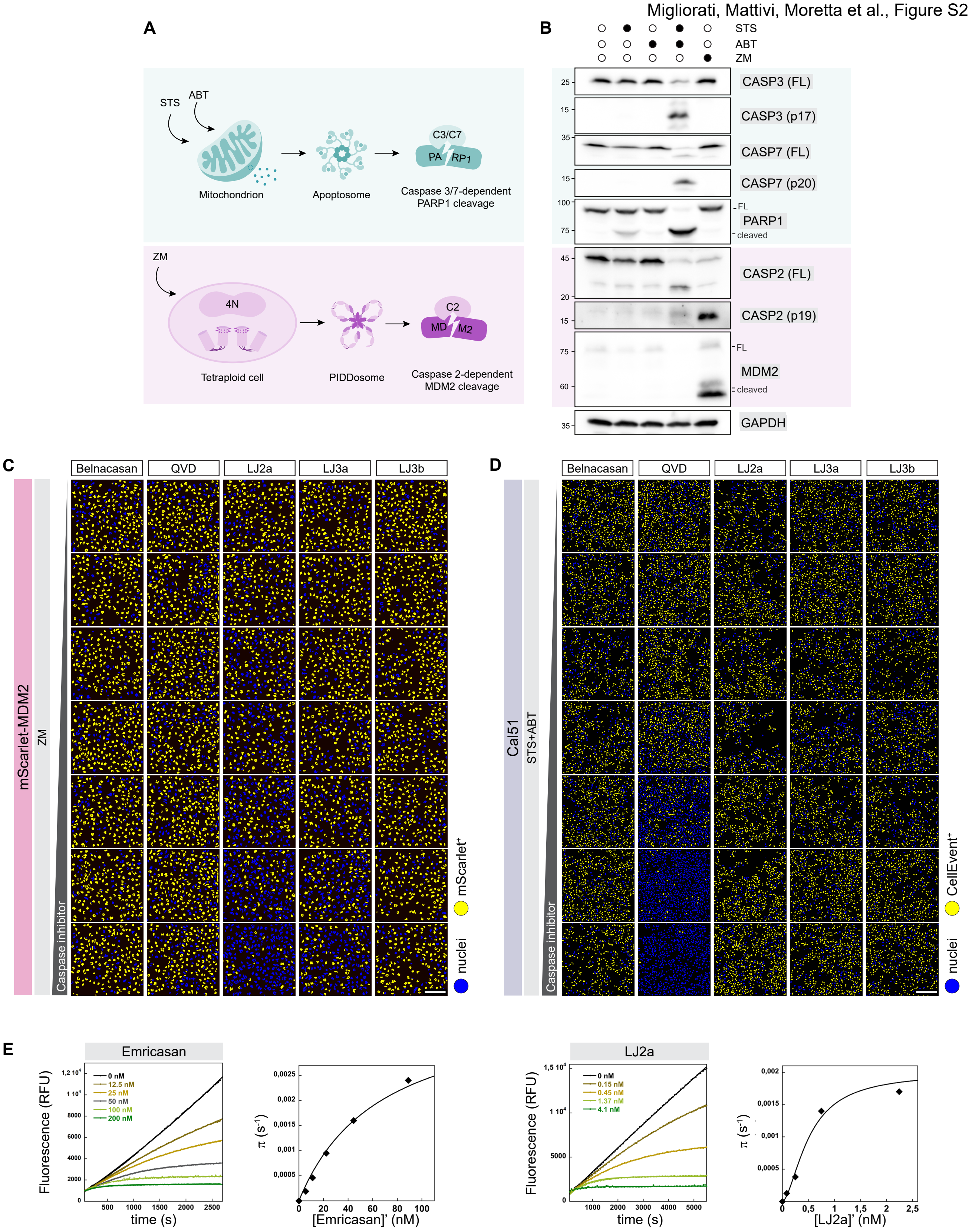


**F**

| Compound | $k_{inact}$ / $K_I$ ($M^{-1}·s^{-1}$) CASP2 | CASP3 | Selectivity index CASP2 / CASP3 |
|---|---|---|---|
| Emricasan | 90 000 | 36 500 | ~ 2.5 |
| QVD | 700 | 167 000 | ~ 0.004 |
| LJ2a | 5 400 000 | 76 000 | ~ 71 |

**Figure S2**. **Representative images from caspase inhibitor profiling screen, related to Figure 2.**
**(A)** Schematic illustration of the cellular pathways engaged by the pharmacological stimuli used in the caspase inhibitor profiling assays. Staurosporine (STS) and ABT-737 (ABT) act on the mitochondrial outer membrane by directly or indirectly modulating members of the BCL-2 protein family, thereby promoting mitochondrial outer membrane permeabilization and downstream effector caspase activation. ZM treatment induces whole-genome doubling, leading to the formation of supernumerary centrosomes.
**(B)** Immunoblot analysis of Cal51 cells treated with vehicle (DMSO), STS, ABT, ZM, or the combined STS + ABT treatment. MDM2, PARP1, and caspases-2, -3, and -7 show distinct cleavage patterns under the different treatment conditions, with ZM treatment associated with Caspase-2 processing and STS + ABT treatment associated with robust effector caspase activation.
**(C)** Representative high-content images of $Cal51^{mScarlet\text{-}MDM2}$ cells treated with ZM in the presence of increasing amounts of the indicated caspase inhibitors. Nuclear mScarlet fluorescence is displayed as binary labeling, with nuclei exceeding the fluorescence threshold pseudo-colored in yellow and nuclei below threshold shown in blue. Images are representative of 3 biological replicates. Scale bar: 200 μm.
**(D)** Representative high-content images of Cal51 cells treated with the combined pro-apoptotic stimulus (STS + ABT) in the presence of increasing amounts of the indicated caspase inhibitors. Effector caspase activity was assessed using the CellEvent reporter and is displayed as binary nuclear labeling, with CellEvent-positive nuclei shown in yellow and negative nuclei in blue. Images are representative of 3 biological replicates. Scale bar: 200 μm.
**(E)** Representative progress curves of Caspase-2 inactivation by Emricasan and LJ2a measured using a continuous fluorometric assay[1]. Caspase-2 (0.1 nM) activity toward Ac-VDVAD-AMC substrate (25 μM, approximately $K_M$) was monitored in the presence of increasing inhibitor concentrations. Release of the fluorescent AMC product was recorded in activity buffer, and data were fitted using a mono-exponential equation as previously described[2] to derive $k_{obs}$ values. Extracted $k_{obs}$ values were plotted against inhibitor concentration and fitted to a hyperbolic equation to determine $k_{inact}$ and $K_I$ values parameters.
**(F)** Table summarizing mean $k_{inact}$ /$K_I$ values for Caspase-2 and Caspase-3 for the indicated inhibitors (Emricasan, QVD, LJ2a), derived from two independent experiments. The selectivity index for Caspase-2 over Caspase-3 corresponds to the ratio of their respective $k_{inact}$ /$K_I$ values for each compound.

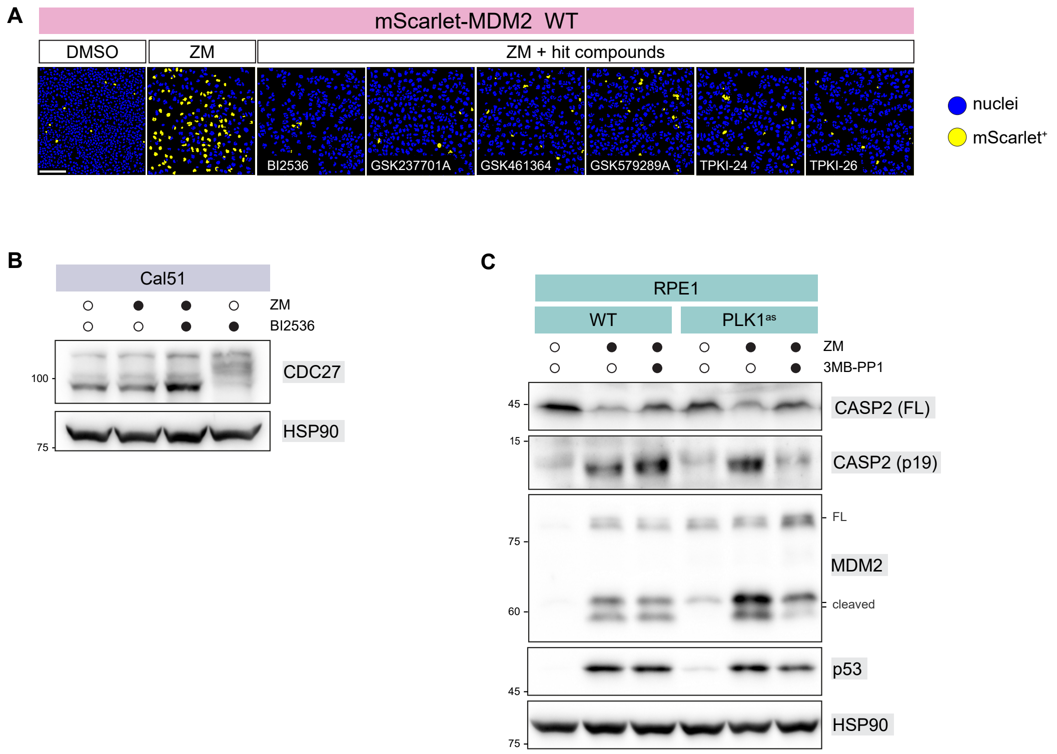


**Figure S3. Supplementary data supporting the identification of PLK1 as a regulator of Caspase-2 activation, related to Figure 3.**
**(A)** Representative images from the high-content screen displaying Cal51$^{mScarlet\text{-}MDM2}$ cells treated with DMSO, ZM, or ZM in combination with each of the six hit compounds (1 µM). Nuclei exceeding the fluorescence threshold are pseudo-colored in yellow, whereas nuclei below threshold are shown in blue. Scale bar: 200 µm.
**(B)** Immunoblot analysis of CDC27 mobility in Cal51 cells treated with DMSO, ZM, BI2536, or ZM + BI2536. BI2536 alone induces a phosphorylation-dependent mobility shift of CDC27, consistent with mitotic arrest, whereas this shift is not observed upon ZM or ZM + BI2536 treatment, consistent with the lack of sustained mitotic arrest under polyploidizing conditions. The first three samples analyzed correspond to those displayed in Fig. 3C.
**(C)** RPE1 cells expressing an analog-sensitive PLK1 allele (PLK1$^{as}$) were treated with ZM in the presence or absence of the ATP analog 3MB-PP1. Acute inhibition of PLK1 results in blunted MDM2 cleavage, phenocopying key aspects of PLK1 inhibition by BI2536 and confirming the on-target requirement for PLK1 activity.

Migliorati, Mattivi, Moretta et al., Figure S4

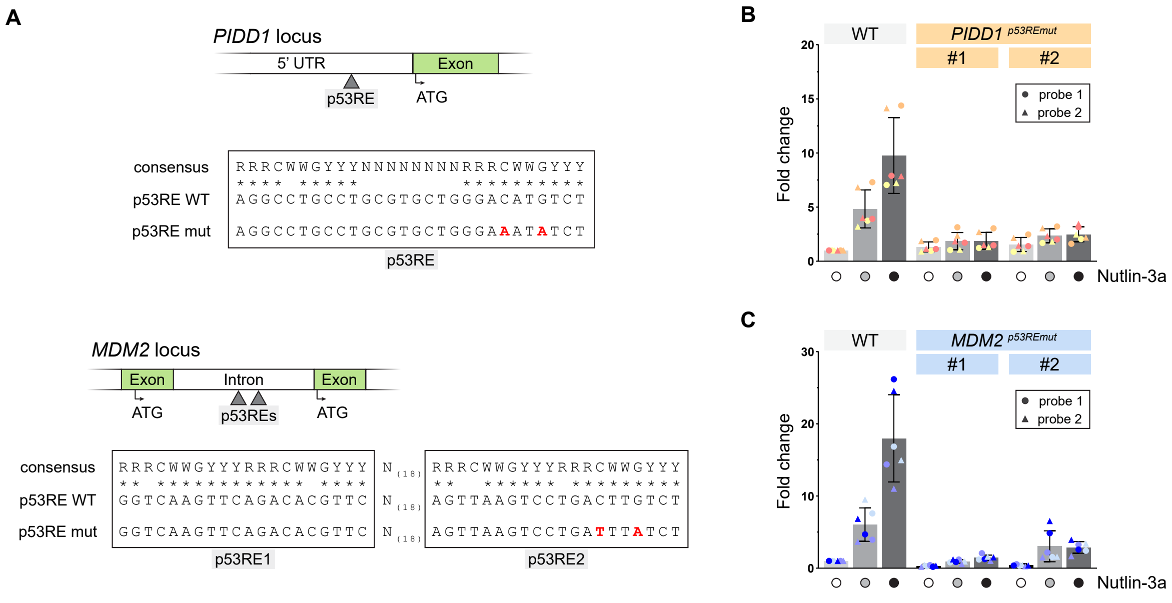


**Figure S4. Validation of p53-dependent transcriptional regulation of *PIDD1* and *MDM2*, related to Figure 4.**

**(A)** Schematic representation of the p53 response elements (p53REs) within the *PIDD1* and *MDM2* promoter regions and the CRISPR-mediated knock-in point mutations introduced to disrupt p53 binding.

**(B)** qPCR analysis of *PIDD1* mRNA levels in wild-type Cal51 cells (WT) and in two Cal51 clones harboring CRISPR-mediated knock-in mutations disrupting the p53 response element in the *PIDD1* promoter (*PIDD1*$^{p53REmut}$) following treatment with the p53 activator Nutlin-3a at two different concentrations (3.3 and 10 µM). Expression levels are shown relative to the vehicle-treated control.

**(C)** qPCR analysis of *MDM2* mRNA levels in wild-type Cal51 cells and in two Cal51 clones harboring CRISPR-mediated knock-in mutations disrupting the p53 response element in the *MDM2* promoter (*MDM2*$^{p53REmut}$) following Nutlin-3a treatment (3.3 and 10 µM). Expression levels are shown relative to the vehicle-treated control.

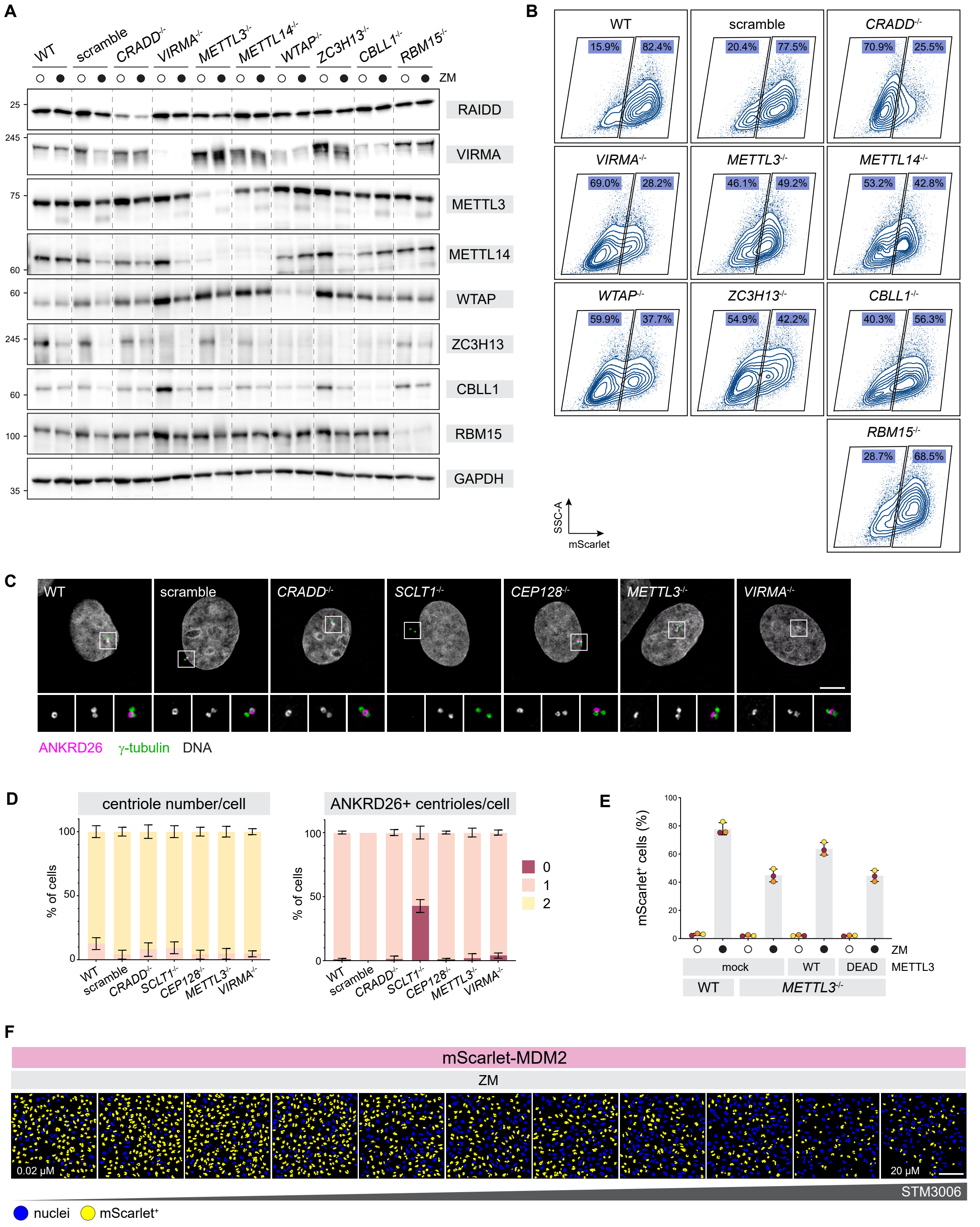


**Figure S5. Supplementary data supporting the involvement of the m6A writer complex in p53-MDM2 pathway output, related to Figure 5.**

**(A)** Immunoblot analysis validating knockout efficiency for each of the seven m6A writer complex components in Cal51 cells shown in Fig. 5C. *CRADD* (RAIDD) knockout is included as a positive control for suppression of pathway-associated readouts. The same loading control (GAPDH) as in Fig. 5C is shown.

**(B)** Representative flow cytometry intensity plots corresponding to the quantification of nuclear mScarlet fluorescence shown in Fig. 5D, illustrating the gating strategy used to define mScarlet-positive and mScarlet-negative cell populations.

**(C)** Immunofluorescence analysis of ANKRD26 localization in Cal51 cells following knockout of the indicated m6A writer complex components. ANKRD26 centrosomal localization is preserved across all perturbations. Magnified insets of boxed regions (without the DNA channel) are shown. Scale bar: 5 μm.

**(D)** Visual scoring of centriole number (left panel) and localization of the distal appendage-associated marker ANKRD26 (right panel) based on immunofluorescence images shown in (C). For each condition, centriole number was scored per cell using γ-tubulin (0, 1, or 2 centrioles per cell) and ANKRD26 localization was scored as present or absent (0 or 1 per cell). Scoring was performed across three biological replicates (n = 50 cells per replicate; 150 cells total).

**(E)** Flow cytometry analysis of mScarlet fluorescence in Cal51$^{mScarlet\text{-}MDM2}$ wild-type (WT), *METTL3*$^{-/-}$, and *METTL3*$^{-/-}$ cells reconstituted with either wild-type (WT) or catalytic-dead (DPPW→APPA) METTL3, following ZM treatment. Bars represent mean ± standard deviation (N = 3 independent biological replicates).

**(F)** Representative high-content imaging data of Cal51$^{mScarlet\text{-}MDM2}$ cells treated with ZM in the presence of increasing amounts of STM3006, corresponding to the reporter-based assays quantified in Fig. 5D. Nuclei exceeding the fluorescence threshold are pseudo-colored in yellow, whereas nuclei below threshold are shown in blue. Scale bar: 200 μm.

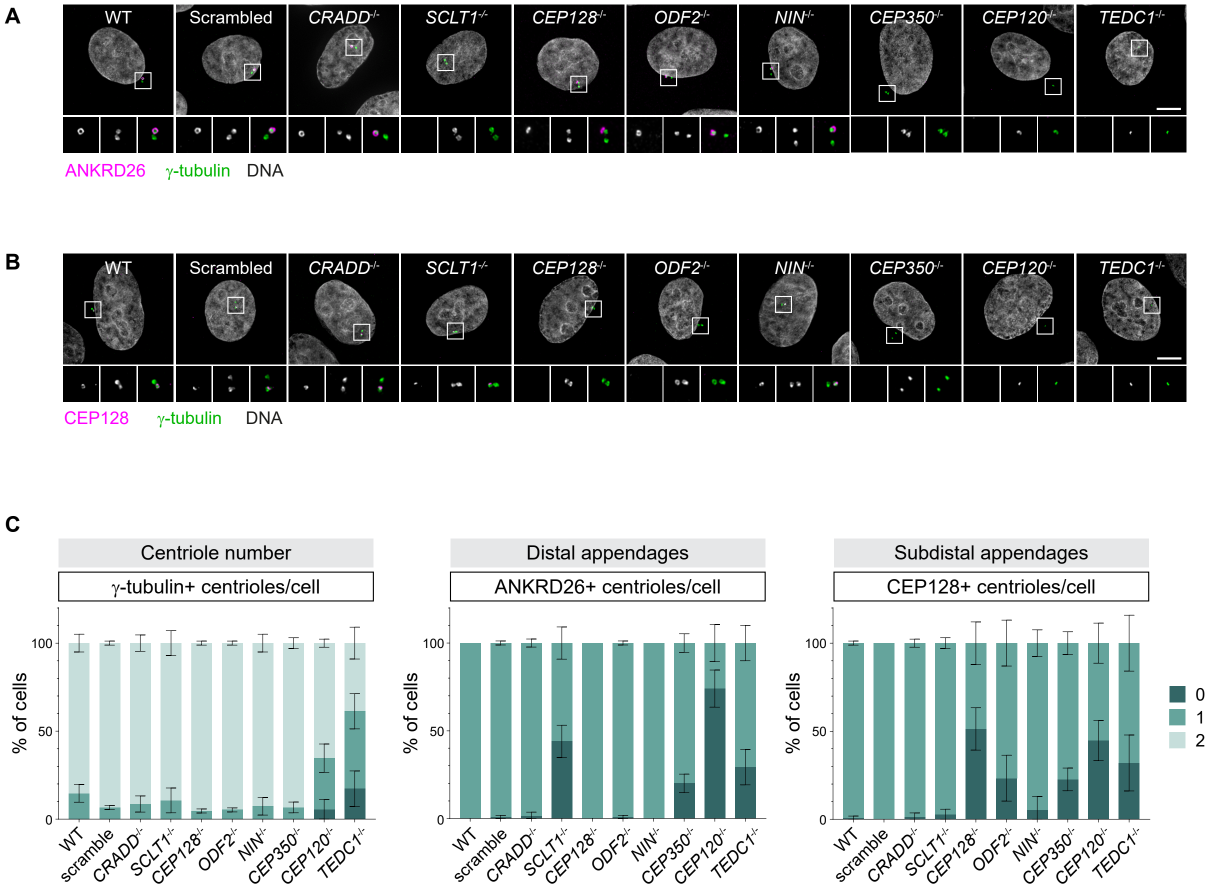


**Figure S6. Supplementary analyses of centrosome integrity and appendage organization, related to Figure 6.**

**(A)** Immunofluorescence analysis of centrosomes in Cal51 cells following acute disruption of the indicated centrosomal components. γ-tubulin marks the pericentriolar material and ANKRD26 marks the centriolar docking site for the PIDDosome (i.e., distal appendages). Magnified insets of boxed regions (without the DNA channel) are shown. Scale bar: 5 μm.

**(B)** Immunofluorescence analysis of subdistal appendages in Cal51 cells following acute disruption of the indicated centrosomal components. γ-tubulin marks the pericentriolar material and CEP128 marks subdistal appendages. Magnified insets of boxed regions (without the DNA channel) are shown. Scale bar: 5 μm.

**(C)** Visual scoring of centrosome number and appendage-associated markers based on immunofluorescence images shown in (A) and (B). For each condition, centrosome number was scored per cell using γ-tubulin (0, 1, or 2 centrioles per cell); ANKRD26 and CEP128 localization were scored as present or absent (0 or 1 per cell). For subdistal appendage assessment, CEP128 was used as the structural marker; accordingly, disruption of NIN (Ninein), which acts downstream of CEP128 in subdistal appendage organization, does not result in loss of CEP128 localization[3]. Scoring was performed independently across three biological replicates, with n = 50 cells scored per replicate (150 cells in total).

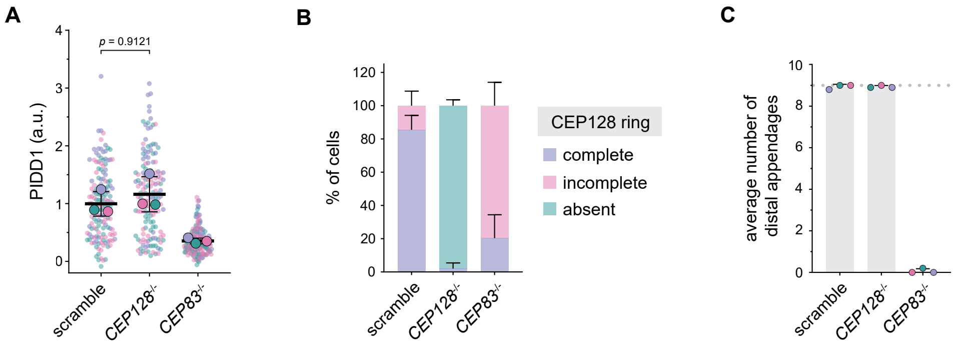


**Figure S7. Supplementary analyses of PIDD1 localization and appendage organization, related to Figure 7.**

**(A)** Superplot of centrosomal PIDD1 fluorescence intensity in RPE1 cells of the indicated genotypes following ZM treatment, from images as in Fig. 7B. n = 150 cells per condition (50 cells per each of 3 biological replicates). Each small dot represents a single cell measurement; same-symbol colors distinguish biological replicates. The mean of each biological replicate is reported (larger dots) ± standard deviation; Kruskal-Wallis test

**(B)** Qualitative analysis of subdistal appendage organization assessed by ultrastructure expansion microscopy (U-ExM) of CEP128 staining in RPE1 cells of the indicated genotypes. In wild-type cells (scramble), CEP128 forms a continuous ring surrounding the centriole. This ring is absent in *CEP128*$^{-/-}$ cells, whereas in *CEP83*$^{-/-}$ cells CEP128 frequently displays partial or discontinuous ring patterns. Qualitative classification was performed across three biological replicates, with n ≥ 15 cells scored per replicate.

**(C)** Quantification of distal appendage number based on U-ExM analysis of CEP83 staining in RPE1 scramble, *CEP128*$^{-/-}$, and *CEP83*$^{-/-}$ cells, from images as in Fig 7C. The number of CEP83-positive appendage structures per centriole was scored, confirming preservation of 9 distal appendages (dotted line) in subdistal appendage-deficient cells. Quantification was performed across three biological replicates, with n ≥ 15 cells scored per replicate.

### Supplementary Movie 1, legend

Representative time-lapse images of asynchronously growing Cal51$^{mScarlet\text{-}MDM2}$ cells treated with vehicle (DMSO, left panels) or ZM (right panels). mScarlet fluorescence is shown in yellow and DNA in red. Time is indicated in minutes. Scale bar: 20 µm.

### Supplementary Table S1

| REAGENT or RESOURCE | SOURCE | IDENTIFIER |
|---|---|---|
| Antibodies | | |
| Rabbit polyclonal anti-CEP83 | Atlas Antibodies | Cat# HPA038161, RRID:AB_10674547 |
| Mouse monoclonal anti-Acetylated Tubulin [clone 6-11B-1] | Sigma-Aldrich | Cat# T7451, RRID:AB_609894 |
| Rabbit polyclonal anti-ANKRD26 | GeneTex | Cat# GTX128255, RRID:AB_2885741 |
| Mouse monoclonal anti-γ-tubulin [clone TU-30] | Thermo Fisher Scientific | Cat# MA1-19421, RRID:AB_1075282 |
| Rabbit polyclonal anti-CEP128 | Atlas antibodies | Cat# HPA001116, RRID:AB_1078323 |
| Mouse monoclonal anti-PIDD1 [clone Anto-1] | Boster Bio | Cat# M10708, RRID:N/A |
| Rabbit polyclonal anti-Centrin 1 | Proteintech | Cat# 12794-1-AP, RRID:AB_2077371 |
| Rat monoclonal anti-Caspase-2 [clone 11B4] | Andreas Strasser, Walter and Eliza Hall Institute of Medical Research | N/A |
| Mouse monoclonal anti-MDM2 [clone IF2] | Thermo Fisher Scientific | Cat# MA1-113, RRID:AB_2536824 |
| Rabbit monoclonal anti-V5-tag [clone D3H8Q] | Cell Signaling Technology | Cat# 13202, RRID:AB_2687461 |
| Rabbit polyclonal anti-p53 | Cell Signaling Technology | Cat# 9282, RRID:AB_331476 |
| Mouse monoclonal anti-MCL1 [clone 22] | Santa Cruz Biotechnology | Cat# sc-12756, RRID:AB_627915 |
| Rabbit polyclonal anti-PARP1 | Cell Signaling Technology | Cat# 9542, RRID:AB_2160739 |
| Rabbit polyclonal anti-Caspase-3 | Cell Signaling Technology | Cat# 9662, RRID:AB_331439 |
| Rabbit polyclonal anti-Caspase-7 | Cell Signaling Technology | Cat# 9492, RRID:AB_2228313 |
| Mouse monoclonal anti-CDC27 [clone 35/CDC27] | BD Biosciences | Cat# 610455, RRID:AB_397828 |
| Mouse monoclonal anti-p21 [clone 2G12] | BD Biosciences | Cat# 564262, RRID:AB_395331 |
| Rabbit monoclonal anti-METTL3 [clone D2I6O] | Cell Signaling Technology | Cat# 96391, RRID:AB_2800261 |
| Rabbit polyclonal anti-METTL14 | Proteintech | Cat# 26158-1-AP, RRID:AB_2800447 |
| Rabbit polyclonal anti-CRADD/RAIDD | Proteintech | Cat# 10401-1-AP, RRID:AB_2085477 |
| Rabbit monoclonal anti-VIRMA [clone D4N8B] | Cell Signaling Technology | Cat# 88358, RRID:AB_2800121 |

| | | |
|---|---|---|
| Rabbit polyclonal anti-WTAP | Cell Signaling Technology | Cat# 56501, RRID:AB_2799512 |
| Rabbit polyclonal anti-ZC3H13 | Boster Bio | Cat# A11022-1, RRID:N/A |
| Rabbit polyclonal anti-CBLL1 | Bethyl | Cat# A302-968A, RRID:AB_10754975 |
| Mouse monoclonal anti-RBM15 [clone 4A1A4] | Proteintech | Cat# 66059-1-Ig, RRID:AB_11042326 |
| Mouse monoclonal anti-HSP90 [clone F-8] | Santa Cruz Biotechnology | Cat# sc-13119, RRID:AB_675659 |
| Rabbit monoclonal anti-GAPDH [clone 14C10] | Cell Signaling Technology | Cat# 2118, RRID:AB_561053 |
| Polyclonal Goat anti-Rat Immunoglobulins/HRP | Thermo Fisher Scientific | Cat# 31470, RRID:AB_228356 |
| Polyclonal Goat Anti-Rabbit Immunoglobulins/HRP | Agilent | Cat# P0448, RRID:AB_2617138 |
| Polyclonal Rabbit Anti-Mouse Immunoglobulins/HRP | Agilent | Cat# P0161, RRID:AB_2687969 |
| Goat polyclonal anti-Mouse IgG (H+L) Highly Cross-Adsorbed Secondary Antibody - Alexa Fluor 488 | Thermo Fisher Scientific | Cat# A-11029, RRID:AB_2534088 |
| Goat polyclonal anti-Mouse IgG (H+L) Highly Cross-Adsorbed Secondary Antibody - Alexa Fluor 555 | Thermo Fisher Scientific | Cat# A-21424, RRID:AB_141780 |
| Goat polyclonal anti-Rabbit IgG (H+L) Highly Cross-Adsorbed Secondary Antibody - Alexa Fluor 488 | Thermo Fisher Scientific | Cat# A-11034, RRID:AB_2576217 |
| Goat polyclonal anti-Rabbit IgG (H+L) Highly Cross-Adsorbed Secondary Antibody - Alexa Fluor 555 | Thermo Fisher Scientific | Cat# A-21429, RRID:AB_2535850 |
| Mouse monoclonal anti-m6A | Synaptic Systems Antibodies | Cat# 202-111 |
| Bacterial and virus strains | | |
| DH5α Competent Cells | Thermo Fisher Scientific | Cat# EC0112 |
| One Shot™ Stbl3™ Chemically Competent E. coli | Thermo Fisher Scientific | Cat# C737303 |
| Chemicals, peptides, and recombinant proteins | | |
| ZM-447439 | MedChemExpress | Cat# HY-10128 |
| Cycloheximide | Thermo Scientific Chemicals | Cat# 357420010 |
| Staurosporine | MedChemExpress | Cat# HY-15141 |
| ABT-737 | MedChemExpress | Cat# HY-50907 |
| BI 2536 | MedChemExpress | Cat# HY-50698 |
| Nutlin-3a | MedChemExpress | Cat# HY-10029 |
| 3MB-PP1 | Cayman Chemical | Cat# 17860 |
| Belnacasan | MedChemExpress | Cat# HY-13205 |
| Emricasan | MedChemExpress | Cat# HY-10396 |
| Q-VD-OPh | MedChemExpress | Cat# HY-12305 |
| LJ2a | Etienne Jacotot, Paris Cité University / Inserm | N/A |
| LJ3a | Etienne Jacotot, Paris Cité University / Inserm | N/A |
| LJ3b | Etienne Jacotot, Paris Cité University / Inserm | N/A |
| STM2457 | Cayman Chemical | Cat# 34280 |
| UZH2 | TargetMol | Cat# T40357 |
| STC-15 | MedChemExpress | Cat# HY-156677 |
| STM3006 | MedChemExpress | Cat# HY-156773 |

| | | |
|---|---|---|
| Doxycycline hyclate | Thermo Scientific Chemicals | Cat# 446060050 |
| Puromycin dihydrochloride | Invivogen | Cat# ant-pr |
| Blasticidin hydrochloride | Invivogen | Cat# ant-bl |
| G418 | Invivogen | Cat# ant-gn |
| NU7441 | Selleckchem | Cat# S2638 |
| Alt-R Cas9 Electroporation Enhancer | IDT | Cat# 1075916 |
| Hoechst 33342 | Thermo Fisher Scientific | Cat# 382065 |
| ProLong™ Gold Antifade Mountant | Thermo Fisher Scientific | Cat# P10144 |
| Hexadimethrine bromide | Sigma-Aldrich | Cat# H9268 |
| DMEM | Thermo Fisher Scientific | Cat# 11960-044 |
| DMEM/F12 1:1 | Thermo Fisher Scientific | Cat# 11320-074 |
| DMEM without phenol red | Thermo Fisher Scientific | Cat# 21063029 |
| RPMI 1640 medium | Thermo Fisher Scientific | Cat# 31870-025 |
| Fetal bovine serum | Thermo Fisher Scientific | Cat# 10270-106 |
| L-glutamine | Thermo Fisher Scientific | Cat# 25030-024 |
| Penicillin-Streptomycin solution | Thermo Fisher Scientific | Cat# 15140-122 |
| Dimethyl sulfoxide | Sigma-Aldrich | Cat# 472301 |
| Human active recombinant caspase-2 | Enzo Life Sciences | Cat# ALX-201-057 |
| Human active recombinant caspase-3 | R&D Systems | Cat# 707-C3 |
| Ac-VDVAD-AMC | Enzo Life Sciences | Cat# ALX-260-060 |
| Ac-DEVD-AMC | Enzo Life Sciences | Cat# ALX-260-031 |
| KCGS Library (v1.0) | Ximbio | Cat# 157681 |
| Formaldehyde | Thermo Fisher Scientific | Cat# BP531-500 |
| Acrylamide Solution | Thermo Fisher Scientific | Cat# BP1402-1 |
| Bis-Acrylamide Solution | Thermo Fisher Scientific | Cat# BP1404-250 |
| Sodium acrylate | Santa Cruz Biotechnology | Cat# SC-236893 |
| TEMED | Sigma-Aldrich | Cat# T9281 |
| Ammonium persulfate | Thermo Fisher Scientific | Cat# BP179-100 |
| Albumin fraction V (from bovine serum) | Merck | Cat# 112018 |
| Critical commercial assays | | |
| CellEvent™ Caspase-3/7 Detection Reagent, green | Thermo Fisher Scientific | Cat# C10423 |
| P3 Primary Cell 4D-Nucleofector® X Kit S | Lonza Bioscience | Cat# V4XP-3032 |
| SE Cell Line 4D-Nucleofector® X Kit S | Lonza Bioscience | Cat# V4XC-1032 |
| NucleoSpin Tissue kit | Macherey-Nagel | Cat# 740952 |
| NucleoSpin Plasmid kit | Macherey-Nagel | Cat# 740588 |
| CompactPrep Plasmid Kits | Qiagen | Cat# 12863 |
| NucleoSpin Gel and PCR Clean-up kit | Macherey-Nagel | Cat# 740609 |
| NucleoSpin RNA Plus kit | Macherey-Nagel | Cat# 740984 |
| RevertAid First Strand cDNA Synthesis Kit | Thermo Fisher Scientific | Cat# K1622 |
| 2x qPCRBIO Probe Mix No-ROX | PCR Biosystems | Cat# PB20.23 |
| Phusion™ High-Fidelity DNA Polymerase | Thermo Fisher Scientific | Cat# F530 |
| Mix2Seq Kit NXP | Eurofins | Cat# 3094-0ONMSK |
| Pierce™ BCA Protein Assay Kit | Thermo Fisher Scientific | Cat# 23225 |
| Amersham™ ECL Select™ Western Blotting Detection Reagent | Cytiva | Cat# RPN2235 |
| m6A RNA Methylation Assay Kit (Colorimetric) | Abcam | Cat# ab185912 |
| PIDD1 Taqman probe (exons 5-6) | IDT | Cat# Hs.PT.58.1440761.g |

| | | |
|---|---|---|
| PIDD1 Taqman probe (exons 10-11) | IDT | Cat# Hs.PT.58.3199598.gs |
| MDM2 Taqman probe (exons 9-11) | IDT | Cat# Hs.PT.58.358457 |
| MDM2 Taqman probe (exons 7-8) | IDT | Cat# Hs.PT.58.4602457 |
| p53 Taqman probe (exons 5-6) | IDT | Cat# Hs.PT.58.38763224.g |
| β-actin Taqman probe (exons 1-2) | IDT | Cat# Hs.PT.39a.22214847 |
| Dynabeads™ Oligo (dT)25 | Invitrogen | Cat# 61005 |
| Metafectene | Biontex | Cat# T020-2.0 |
| SiR-DNA kit | Spirochrome | Cat# SC007 |
| Experimental models: Cell lines | | |
| CL01: Cal51 | Yossi Shiloh (Tel Aviv University) | N/A |
| CL02: hTERT-RPE1 | Stephan Geley (Medical University of Innsbruck) | N/A |
| CL03: A549 | ATCC | CCL-185 |
| CL04: U2OS | Anna Cereseto (University of Trento) | N/A |
| CL05: HCT116 | Alessandra Bisio (University of Trento) | N/A |
| CL06: MCF7 | DSMZ | ACC 115 |
| CL07: Nalm6 | DSMZ | ACC 128 |
| CL08: HepG2 | Graziano Lolli (University of Trento) | N/A |
| CL09: Cal51$^{mScarlet\text{-}MDM2}$ (clone 1) | This paper | N/A |
| CL10: Cal51$^{mScarlet\text{-}MDM2}$ (clone 2) | This paper | N/A |
| CL11: Cal51$^{mScarlet\text{-}MDM2}$ *CEP83*$^{-/-}$ | This paper | N/A |
| CL12: Cal51 *TP53*$^{-/-}$ | This paper | N/A |
| CL13: hTERT-RPE1 *TP53*$^{-/-}$ (clone 1) | This paper | N/A |
| CL14: hTERT-RPE1 *TP53*$^{-/-}$ (clone 2) | This paper | N/A |
| CL15: hTERT-RPE1 p53 R175H (clone 1) | This paper | N/A |
| CL16: hTERT-RPE1 p53 R175H (clone 2) | This paper | N/A |
| CL17: hTERT-RPE1 p53 R248Q (clone 1) | This paper | N/A |
| CL18: hTERT-RPE1 p53 R248Q (clone 2) | This paper | N/A |
| CL19: Cal51 *MDM2* $^{\Delta p53RE}$ (clone 1) | This paper | N/A |
| CL20: Cal51 *MDM2* $^{\Delta p53RE}$ (clone 2) | This paper | N/A |
| CL21: Cal51 *PIDD1* $^{\Delta p53RE}$ (clone 1) | This paper | N/A |
| CL22: Cal51 *PIDD1* $^{\Delta p53RE}$ (clone 2) | This paper | N/A |
| CL23: Cal51 rtTR | This paper | N/A |
| CL24: Cal51 Cal51$^{mScarlet\text{-}MDM2}$ (clone 1) rtTR | This paper | N/A |
| CL25: hTERT-RPE1 PLK1$^{as}$ | Prasad Jallepalli (Memorial Sloan Kettering Cancer Center) | N/A |
| CL26: HEK 293T | Ulrich Maurer (University of Freiburg) | N/A |
| CL27: hTERT-RPE1 *CEP83*$^{-/-}$ (lentiCRISPR-V2) | This paper | N/A |
| CL28: hTERT-RPE1 *CEP128*$^{-/-}$ (lentiCRISPR-V2) | This paper | N/A |
| Oligonucleotides | | |

| | | |
|---|---|---|
| crRNA: MDM2 (reporter generation)<br>GTATTGCACATTTGCCTACA | IDT | N/A |
| Primer: MDM2 reporter cassette (FW)<br>TTTCCAGTTTTCATCGTGTCTTTTTTTTCCTTGTAGGCAAACCATGGCTGAACAAGATGG | Metabion | N/A |
| Primer: MDM2 reporter cassette (RV)<br>CAGCACCATCAGTAGGTACAGACATGTTGGTATTGCACATGCTTCCACTACCCTTGTACA | Metabion | N/A |
| Primer: mScarlet-MDM2 genotyping (FW)<br>CGATTGGAGGGTAGACCTGTG | Metabion | N/A |
| Primer: mScarlet-MDM2 genotyping (RV)<br>TTTTAACTCCACGCAGTTACGC | Metabion | N/A |
| crRNA: *PIDD1*$^{\Delta p53RE}$<br>CCTGCGTGCTGGGACATGTC | IDT | N/A |
| Ultramer: *PIDD1*$^{\Delta p53RE}$<br>A*C*CGTTGCAGCCATCGCCCACCGACGGTCCTTGGAGGCCAGATATTTCCCAGCACGCAGGCAGGCCTGTCCAGGCAGCGCCCGGGG*A*A | IDT | N/A |
| Primer: *PIDD1*$^{\Delta p53RE}$ genotyping (FW)<br>GACCAGGACTGAAGCCTCAC | Metabion | N/A |
| Primer: *PIDD1*$^{\Delta p53RE}$ genotyping (RV)<br>GCAGAGCGAGAGATGGAGAC | Metabion | N/A |
| crRNA: *MDM2*$^{\Delta p53RE}$<br>TCCTGACTTGTCTCCAGCTG | IDT | N/A |
| Ultramer: *MDM2*$^{\Delta p53RE}$<br>C*A*GACACGTTCCGAAACTGCAGTAAAAGGAGTTAAGTCCTGATTTATCTCCAGCTGGGGCTATTTAAACCATGCATTTTCCCAGCT*G*T | IDT | N/A |
| Primer: *MDM2*$^{\Delta p53RE}$ genotyping (FW)<br>GAGGTCCGGATGATCGCAG | Metabion | N/A |
| Primer: *MDM2*$^{\Delta p53RE}$ genotyping (RV)<br>AACTCCACGCAGTTACGCCA | Metabion | N/A |
| crRNA: *TP53*$^{-/-}$<br>TCCATTGCTTGGGACGGCAA | IDT | N/A |
| Primer: *TP53*$^{-/-}$ genotyping (FW)<br>TTGGCGTCTACACCTCAGGA | Metabion | N/A |
| Primer: *TP53*$^{-/-}$ genotyping (RV)<br>GCAGTCAGATCCTAGCGTCG | Metabion | N/A |
| crRNA: p53 R175H<br>AGCACATGACGGAGGTTGTG | IDT | N/A |
| Ultramer: p53 R175H<br>C*C*ATCTACAAGCAGTCACAGCACATGACGGAGGTTGTCAGACACTGCCCCCACCATGAGCGCTGCTCAGATAGCGATGGTGAG*C*A | IDT | N/A |
| Primer: p53 R175H genotyping (FW)<br>GCACCACCACACTATGTCGA | Metabion | N/A |
| Primer: p53 R175H genotyping (RV)<br>CGCCAACTCTCTCTAGCTCG | Metabion | N/A |
| crRNA: p53 R248Q<br>GCATGGGCGGCATGAACCGG | IDT | N/A |
| Ultramer: p53 R248Q<br>A*A*CTACATGTGTAACAGTTCCTGCATGGGCGGCATGAACCAGAGACCCATCCTCACCATCATCACACTGGAAGACTCCAGGTC*A*G | IDT | N/A |
| Primer: p53 R248Q genotyping (FW)<br>GGAGGGGTTAAGGGTGGTTG | Metabion | N/A |

| | | |
|---|---|---|
| Primer: p53 R248Q genotyping (RV) CAGTAAGGAGATTCCCCGCC | Metabion | N/A |
| crRNA: *CEP83*$^{-/-}$ CTCCTCTTAGTTCTTCAAGC | IDT | N/A |
| Primer: *CEP83*$^{-/-}$ genotyping (FW) AGTTTCATGGCTCCCTTACATGG | Eurofins | N/A |
| Primer: *CEP83*$^{-/-}$ genotyping (RV) CCCCATCTCCCACCAAAGTAAA | Eurofins | N/A |
| Alt-R® CRISPR-Cas9 tracrRNA | IDT | Cat# 1072534 |
| Primer: lentiCRISPR-V2 *mCD8*$^{-/-}$ (FW) caccgGTGTTGGGGTCCGTTTCGCA | Eurofins | N/A |
| Primer: lentiCRISPR-V2 *mCD8*$^{-/-}$ (RV) aaacTGCGAAACGGACCCCAACACc | Eurofins | N/A |
| Primer: lentiCRISPR-V2 *CRADD*$^{-/-}$ (FW) caccgCGCTCACTTCGCCTGGAGCT | Eurofins | N/A |
| Primer: lentiCRISPR-V2 *CRADD*$^{-/-}$ (RV) aaacAGCTCCAGGCGAAGTGAGCGc | Eurofins | N/A |
| Primer: lentiCRISPR-V2 *CEP83*$^{-/-}$ (FW) caccgAAGAATACAGGTGCGGCAGT | Eurofins | N/A |
| Primer: lentiCRISPR-V2 *CEP83*$^{-/-}$ (RV) aaacACTGCCGCACCTGTATTCTTc | Eurofins | N/A |
| Primer: lentiCRISPR-V2 *SCLT1*$^{-/-}$ (FW) caccgACTGAGTATGATAAACACCT | Eurofins | N/A |
| Primer: lentiCRISPR-V2 *SCLT1*$^{-/-}$ (RV) aaacAGGTGTTTATCATACTCAGTc | Eurofins | N/A |
| Primer: lentiCRISPR-V2 *CEP128*$^{-/-}$ (FW) caccgGAATCACTTTGACACATGTG | Eurofins | N/A |
| Primer: lentiCRISPR-V2 *CEP128*$^{-/-}$ (RV) aaacCACATGTGTCAAAGTGATTCc | Eurofins | N/A |
| Primer: lentiCRISPR-V2 *ODF2*$^{-/-}$ (FW) caccgGGCACAGCTTCGGTCCAAAG | Eurofins | N/A |
| Primer: lentiCRISPR-V2 *ODF2*$^{-/-}$ (RV) aaacCTTTGGACCGAAGCTGTGCCc | Eurofins | N/A |
| Primer: lentiCRISPR-V2 *NIN*$^{-/-}$ (FW) caccgCTGGAAGACGCAACGCAGTG | Eurofins | N/A |
| Primer: lentiCRISPR-V2 *NIN*$^{-/-}$ (RV) aaacCACTGCGTTGCGTCTTCCAGc | Eurofins | N/A |
| Primer: lentiCRISPR-V2 *CEP350*$^{-/-}$ (FW) caccgACCATCCGAGTCTATAGGAC | Eurofins | N/A |
| Primer: lentiCRISPR-V2 *CEP350*$^{-/-}$ (RV) aaacGTCCTATAGACTCGGATGGTc | Eurofins | N/A |
| Primer: lentiCRISPR-V2 *CEP120*$^{-/-}$ (FW) caccgGTGGAGCTAGCCCCAACTGT | Eurofins | N/A |
| Primer: lentiCRISPR-V2 *CEP120*$^{-/-}$ (RV) aaacACAGTTGGGGCTAGCTCCACc | Eurofins | N/A |
| Primer: lentiCRISPR-V2 *TEDC1*$^{-/-}$ (FW) caccgGGCACTGGCACAACTACCTG | Eurofins | N/A |
| Primer: lentiCRISPR-V2 *TEDC1*$^{-/-}$ (RV) aaacCAGGTAGTTGTGCCAGTGCCc | Eurofins | N/A |
| Primer: lentiCRISPR-V2 *VIRMA*$^{-/-}$ (FW) caccgGAAGTCCGAGTCATACCCCC | Eurofins | N/A |
| Primer: lentiCRISPR-V2 *VIRMA*$^{-/-}$ (RV) aaacGGGGGTATGACTCGGACTTCc | Eurofins | N/A |
| Primer: lentiCRISPR-V2 *METTL3*$^{-/-}$ (FW) caccgGGACACGTGGAGCTCTATCC | Eurofins | N/A |

| | | |
|---|---|---|
| Primer: lentiCRISPR-V2 *METTL3*$^{-/-}$ (RV) aaacGGATAGAGCTCCACGTGTCCc | Eurofins | N/A |
| Primer: lentiCRISPR-V2 *METTL14*$^{-/-}$ (FW) caccgATAGTACAATTCGACCAGGT | Eurofins | N/A |
| Primer: lentiCRISPR-V2 *METTL14*$^{-/-}$ (RV) aaacACCTGGTCGAATTGTACTATc | Eurofins | N/A |
| Primer: lentiCRISPR-V2 *WTAP*$^{-/-}$ (FW) caccgGAAGTGTCGAATGCTTATCC | Eurofins | N/A |
| Primer: lentiCRISPR-V2 *WTAP*$^{-/-}$ (RV) aaacGGATAAGCATTCGACACTTCc | Eurofins | N/A |
| Primer: lentiCRISPR-V2 *ZC3H13*$^{-/-}$ (FW) caccgTGTCCGGAGAATGTTCACGC | Eurofins | N/A |
| Primer: lentiCRISPR-V2 *ZC3H13*$^{-/-}$ (RV) aaacGCGTGAACATTCTCCGGACAc | Eurofins | N/A |
| Primer: lentiCRISPR-V2 *CBLL1*$^{-/-}$ (FW) caccgGGAGCACGAATATCCTCATG | Eurofins | N/A |
| Primer: lentiCRISPR-V2 *CBLL1*$^{-/-}$ (RV) aaacCATGAGGATATTCGTGCTCCc | Eurofins | N/A |
| Primer: lentiCRISPR-V2 *RMB15*$^{-/-}$ (FW) caccgCTGCGATCTAGGCTATCCAG | Eurofins | N/A |
| Primer: lentiCRISPR-V2 *RMB15*$^{-/-}$ (RV) aaacCTGGATAGCCTAGATCGCAGc | Eurofins | N/A |
| Primer: LentiCRISPR-V2 *CEP83*$^{-/-}$ genotyping (FW) TTAAGACATTTCAGATAGGTGACTTGACTCCC | Eurofins | N/A |
| Primer: LentiCRISPR-V2 *CEP83*$^{-/-}$ genotyping (RV) GGAAAAGCTTTGGAGTAGTCAGAGTTTTGG | Eurofins | N/A |
| Primer: LentiCRISPR-V2 *CEP128*$^{-/-}$ genotyping (FW) GCACATTGTTCGCTTCAAAAGTG | Eurofins | N/A |
| Primer: LentiCRISPR-V2 *CEP128*$^{-/-}$ genotyping (RV) CAGATTGTTTCTCCAACGCTCG | Eurofins | N/A |
| Primer: barcoded forward NGS primer AATGATACGGCGACCACCGAGATCTACACTCTTTCCCTACACGACGCTCTTCCGATCTn[1-8]GTGGAAAGGACGAAACACCG | Eurofins | N/A |
| Primer: barcoded reverse NGS primer #1 CAAGCAGAAGACGGCATACGAGATacacgatcGTGACTGGAGTTCAGACGTGTGCTCTTCCGATCTn[1-8]CCAATTCCCACTCCTTTCAAGACCT | Eurofins | N/A |
| Primer: barcoded reverse NGS primer #2 CAAGCAGAAGACGGCATACGAGATcgcgcggtGTGACTGGAGTTCAGACGTGTGCTCTTCCGATCTn[1-8]CCAATTCCCACTCCTTTCAAGACCT | Eurofins | N/A |
| Primer: barcoded reverse NGS primer #3 CAAGCAGAAGACGGCATACGAGATcatgatcgGTGACTGGAGTTCAGACGTGTGCTCTTCCGATCTn[1-8]CCAATTCCCACTCCTTTCAAGACCT | Eurofins | N/A |
| Primer: barcoded reverse NGS primer #4 CAAGCAGAAGACGGCATACGAGATcgttaccaGTGACTGGAGTTCAGACGTGTGCTCTTCCGATCTn[1-8]CCAATTCCCACTCCTTTCAAGACCT | Eurofins | N/A |
| Primer: barcoded reverse NGS primer #5 CAAGCAGAAGACGGCATACGAGATatgcgcagGTGACTGGAGTTCAGACGTGTGCTCTTCCGATCTn[1-8]CCAATTCCCACTCCTTTCAAGACCT | Eurofins | N/A |

| | | |
|---|---|---|
| Primer: barcoded reverse NGS primer #6 CAAGCAGAAGACGGCATACGAGATtagcgctcGTGACTGGAGTTCAGACGTGTGCTCTTCCGATCTn[1-8]CCAATTCCCACTCCTTTCAAGACCT | Eurofins | N/A |
| Primer: barcoded reverse NGS primer #7 CAAGCAGAAGACGGCATACGAGATactgagcgGTGACTGGAGTTCAGACGTGTGCTCTTCCGATCTn[1-8]CCAATTCCCACTCCTTTCAAGACCT | Eurofins | N/A |
| Primer: mutagenesis from METTL3 “APPA” to “WT” with SPRINP protocol (FW) GTTTGCAGTTGTGATGGCTGACCCACCCTGGGATATTCACATGGAACTGC | Eurofins | N/A |
| Primer: mutagenesis from METTL3 “APPA” to “WT” with SPRINP protocol (RV) GCAGTTCCATGTGAATATCCCAGGGTGGGTCAGCCATCACAACTGCAAAC | Eurofins | N/A |
| Recombinant DNA | | |
| pcDNA5/FRT/TO | Thermo Fisher Scientific | V652020 |
| pcDNA5/FRT/TO NeoR-T2A-V5-mScarlet | This paper | N/A |
| FUW-tetO-MCS+ | Alessio Zippo (University of Trento) | Modified version of Addgene plasmid #84008 |
| pLenti-ubc-rtTR-M3 | Andreas Villunger | N/A |
| lentiCRISPR v2 | Feng Zhang | Addgene plasmid #52961, RRID:Addgene_52961 |
| pCMV-VSV-G | Bob Weinberg | Addgene plasmid #8484, RRID:Addgene_8454 |
| psPAX2 | Didier Trono | Addgene plasmid #12260, RRID:Addgene_12260 |
| Human CRISPR Knockout Pooled Library (Brunello) | Root, Doench | Addgene #73179 |
| METTL3_cDNA_APPA | This paper | N/A |
| Software and algorithms | | |
| Fiji/ImageJ2 (version 2.16.0) | Schneider et al.[4] | https://imagej.net/software/fiji/downloads |
| FlowJo (version: 11) | BD Bioscience | https://flowjo.com/flowjo/download |
| GraphPad Prism (version 10.0) | GraphPad | https://www.graphpad.com/how-to-buy/ |
| CFX Manager (version 3.1) | Bio-Rad | N/A |
| RStudio (version 2026.01.0; R version 4.5.2) using the clusterProfiler, org.Hs.eg.db, enrichplot, ggplot2, ComplexHeatmap, GO.db, readr, dplyr, enrichplot and patchwork packages | N/A | https://posit.co/downloads/ |
| ICE | Conant et al.[5] | http://ice.synthego.com/ |
| CRISPOR | Concordet and Haeussler[6] | https://crispor.gi.ucsc.edu |
| Cutadapt | Martin | https://doi.org/10.14806/ej.17.1.200 |
| Bowtie2 | Langmead & Salzberg[7] | https://doi.org/10.1038/nmeth.1923 |

| | | |
|---|---|---|
| ReCo | Wegner & Kaulich[8] | https://academic.oup.com/bioinformatics/article/39/8/btad448/7229558 |
| MAGeCK | Li et al.[9] | https://doi.org/10.1186/s13059-014-0554-4 |
| DESeq2 | Love et al.[10] | https://doi.org/10.1186/s13059-014-0550-8 |
| STAR | Dobin et al.[11] | https://doi.org/10.1093/bioinformatics/bts635 |
| Huygens Professional (via Huygens Remote Manager, version 3.10) | Scientific Volume Imaging | N/A |
| NIS Elements (version 5.42.06) | Nikon Instruments Inc | N/A |
| KaleidaGraph | Synergy Software | N/A |
| Other | | |
| 4–20% Criterion™ TGX Stain-Free™ Protein Gel | Bio-Rad | Cat# 5678095 |
| Amersham™ Protran® Western blotting membranes, nitrocellulose | Cytiva | Cat# GE10600001 |
| ECL Select™ Western Blotting Detection Reagent | Cytiva Amersham™ | Cat# 12644055 |
| PhenoPlate, 384-well black optically-clear bottom plates | Revvity | Cat# 6057302 |
| SpectraPlate, 384-well clear, sterile, tissue culture-treated plates | Revvity | Cat# 6007650 |
| clear, sterile, tissue culture-treated 384-well plates | Thermo Fisher Scientific | Cat# 164688 |
| μ-Slide 8 Well | Ibidi | Cat# 80826 |

### Supplementary References


1. Doucet, C., Pochet, L., Thierry, N., Pirotte, B., Delarge, J., and Reboud-Ravaux, M. (1999). 6-Substituted 2-oxo-2H-1-benzopyran-3-carboxylic acid as a core structure for specific inhibitors of human leukocyte elastase. J Med Chem *42*, 4161-4171. 10.1021/jm990070k.
2. Bosc, E., Anastasie, J., Soualmia, F., Coric, P., Kim, J.Y., Wang, L.Q., Lacin, G., Zhao, K., Patel, R., Duplus, E., et al. (2022). Genuine selective caspase-2 inhibition with new irreversible small peptidomimetics. Cell Death Dis *13*, 959. 10.1038/s41419-022-05396-2.
3. Mazo, G., Soplop, N., Wang, W.J., Uryu, K., and Tsou, M.F. (2016). Spatial Control of Primary Ciliogenesis by Subdistal Appendages Alters Sensation-Associated Properties of Cilia. Dev Cell *39*, 424-437. 10.1016/j.devcel.2016.10.006.
4. Schindelin, J., Arganda-Carreras, I., Frise, E., Kaynig, V., Longair, M., Pietzsch, T., Preibisch, S., Rueden, C., Saalfeld, S., Schmid, B., et al. (2012). Fiji: an open-source platform for biological-image analysis. Nat Methods *9*, 676-682. 10.1038/nmeth.2019.
5. Conant, D., Hsiau, T., Rossi, N., Oki, J., Maures, T., Waite, K., Yang, J., Joshi, S., Kelso, R., Holden, K., et al. (2022). Inference of CRISPR Edits from Sanger Trace Data. CRISPR J *5*, 123-130. 10.1089/crispr.2021.0113.

6. Concordet, J.P., and Haeussler, M. (2018). CRISPOR: intuitive guide selection for CRISPR/Cas9 genome editing experiments and screens. Nucleic Acids Res *46*, W242-W245. 10.1093/nar/gky354.
7. Langmead, B., and Salzberg, S.L. (2012). Fast gapped-read alignment with Bowtie 2. Nat Methods *9*, 357-359. 10.1038/nmeth.1923.
8. Wegner, M., and Kaulich, M. (2023). ReCo: automated NGS read-counting of single and combinatorial CRISPR gRNAs. Bioinformatics *39*. 10.1093/bioinformatics/btad448.
9. Li, W., Xu, H., Xiao, T., Cong, L., Love, M.I., Zhang, F., Irizarry, R.A., Liu, J.S., Brown, M., and Liu, X.S. (2014). MAGeCK enables robust identification of essential genes from genome-scale CRISPR/Cas9 knockout screens. Genome Biol *15*, 554. 10.1186/s13059-014-0554-4.
10. Love, M.I., Huber, W., and Anders, S. (2014). Moderated estimation of fold change and dispersion for RNA-seq data with DESeq2. Genome Biol *15*, 550. 10.1186/s13059-014-0550-8.
11. Dobin, A., Davis, C.A., Schlesinger, F., Drenkow, J., Zaleski, C., Jha, S., Batut, P., Chaisson, M., and Gingeras, T.R. (2013). STAR: ultrafast universal RNA-seq aligner. Bioinformatics *29*, 15-21. 10.1093/bioinformatics/bts635.
